# Dynamic Temporal Changes in Urinary Proteome of Pregnant Wild-Type Mice Carrying Heterozygous 3xTg-AD Fetuses

**DOI:** 10.64898/2026.07.23.740247

**Authors:** Lili Guo, Youhe Gao

## Abstract

Alzheimer’s disease (AD) is a progressive neurodegenerative disorder with insidious onset. At present, effective early diagnostic biomarkers are scarce, and few studies have explored ultra-early molecular alterations during embryonic development. Urinary proteomics boasts unique strengths including complete non-invasiveness, repeatable sequential sampling and high detection sensitivity, which provides a potential technical strategy for prenatal monitoring of congenital disorders. In this study, we constructed an experimental group by mating wild-type female mice with 3xTg transgenic male mice to obtain pregnant dams carrying heterozygous AD-susceptible fetuses, while wild-type male-female mating was set as the blank control. Urine samples were consecutively collected at 10 time points from gestational day 1 (D1) to D19. Label-free quantitative proteomics was adopted to screen differentially expressed proteins, and Gene Ontology (GO) enrichment analysis was carried out to interpret temporal biological processes. The results showed that stable intergroup differential proteins could be detected as early as the implantation stage (D1), and differential protein profiles existed throughout the whole gestation period. The quantity and expression trend of differential proteins exhibited obvious temporal dynamics, and permutation tests verified that the intergroup differences were not random noise. Paternally inherited AD-causing mutations could trigger systematic molecular responses in maternal mice at the early embryonic stage. This study for the first time characterized the dynamic urinary proteomic landscape of maternal mice that reflects fetal AD susceptibility. It demonstrates that maternal urine can mirror molecular signatures related to fetal AD development, offering fundamental animal experimental data for subsequent screening of prenatal non-invasive monitoring biomarkers and research on the embryonic origin of AD.

## 1. Introduction

Alzheimer’s disease (AD) is an age-related neurodegenerative disorder clinically characterized by cognitive impairment and progressive dementia^[1]^.Its neuropathological hallmarks include extracellular β-amyloid (Aβ) plaques and intraneuronal neurofibrillary tangles composed of hyperphosphorylated tau protein^[2]^, making it a common cause of acquired cognitive impairment in middle-aged and elderly populations. With the accelerating global population aging, AD has emerged as a major public health challenge worldwide. The number of people affected by AD globally is projected to increase from 50 million in 2010 to 113 million^[3]^. In China, among the population aged 60 years and above, the overall prevalence of dementia is 6.04%, with AD accounting for 3.9%; sporadic AD constitutes more than 95% of all cases. AD and related dementias rank as the fifth leading cause of death in China, and have become one of the most costly, fatal, and burdensome diseases, posing a serious threat to public health and sustainable social development^[4]^. Currently, clinical diagnosis of AD relies largely on mid-to-late-stage neuroimaging alterations and cognitive function assessments^[5–7]^, lacking early diagnostic methods and curative interventions, thereby missing the optimal window for therapeutic intervention. Therefore, the identification of specific biomarkers during the early, asymptomatic stages of AD is critical for achieving early screening, early diagnosis, and early treatment of the disease.

Despite decades of intensive research, the precise pathogenesis of AD remains incompletely elucidated. The classical amyloid-β (Aβ) cascade hypothesis and the traditional aging theory can only partially explain disease progression after adulthood, but fail to account for the substantial inter-individual variability in age of onset and disease risk among individuals carrying the same genotype^[5,7]^. In 1990, the Developmental Origins of Health and Disease (DOHaD) theory first demonstrated that the intrauterine environment can permanently reprogram fetal organ development and lock in susceptibility phenotypes for chronic diseases in adulthood^[8]^. In recent years, multiple birth cohort studies and animal experiments have confirmed that the pathological initiation of AD can be traced back to the intrauterine developmental stage. The window periods for fetal hippocampal and prefrontal cortical neurogenesis and blood-brain barrier development (mid-to-late gestation) have been identified as critical nodes for fetal programming of AD^[9,10]^. Maternal inflammation, metabolic abnormalities, and intrauterine microenvironment disturbances can permanently alter neural stem cell proliferation and differentiation^[11]^ as well as microglial innate immune phenotypes^[12]^. These perturbations, through epigenetic programming, abnormal neural circuit development, and remodeling of brain immune homeostasis, programmatically increase offspring susceptibility to AD in middle and old age^[13,14]^. Collectively, these findings provide theoretical support for the embryonic origin of AD. However, whether non-invasive identification of AD susceptibility status can be achieved during the fetal period has not yet been experimentally demonstrated.

Previous studies on AD biomarkers have mainly focused on brain tissue, cerebrospinal fluid (CSF), peripheral serum and other specimens from affected patients, centering on pathological molecular alterations occurring after disease onset, while rarely investigating preclinical molecular changes during the embryonic origination stage of the disease^[15,16]^. As a non-invasive biofluid that allows continuous and dynamic sampling, urine accurately reflects systemic metabolic and pathological molecular shifts in the body. Compared with blood and CSF, urine is free of interference from high-abundance proteins, facilitating the screening of low-abundance specific differential proteins, and thus serving as an ideal matrix for screening early disease biomarkers^[17,18]^. Accumulating evidence has verified that neurodegenerative disorders trigger aberrant proteomic profiles in peripheral body fluids, and urinary proteomics is capable of capturing covert early signals of nervous system lesions^[19,20]^. Nevertheless, it remains unclear whether pathological alterations in offspring with hereditary AD can induce distinct urinary protein changes in mothers during the early embryonic development period.

The C57BL/6 mouse is a globally standardized inbred strain with a highly consistent genetic background among individuals. Due to this genetic uniformity, phenotypic changes introduced by transgenes or gene knockouts can be directly attributed to the target gene rather than to genetic background or other confounding factors^[21,22]^. The 3xTg (APP/PS1/Tau) mouse is a classic transgenic model of AD. This strain simultaneously integrates three major familial AD causative mutations: the Swedish mutation (APPSwe), the MAPT P301L mutation, and the PSEN1 M146V knock-in mutation. By co-injecting the APP and tau transgenes into single-cell embryos carrying the PS1 knock-in mutation, both transgenes are integrated into the same genomic locus, ensuring genetic background consistency. APP and tau are driven by the neuron-specific Thy1.2 promoter, enabling stable recapitulation of the pathological progression of human AD, including amyloid plaque deposition, neurofibrillary tangles, and cognitive decline. Consequently, this model has been widely used in animal studies investigating AD mechanisms^[23–25]^.

Based on these considerations, to avoid the confounding effects of the maternal transgenic background on gestational pathology and to precisely dissect the maternal urinary molecular profile changes mediated by the fetal AD genetic susceptibility background, we designed two breeding groups in this study. In the experimental group, 3xTg-AD male mice were paired with wild-type female mice, producing offspring heterozygous for the three AD-causing mutations, while the maternal dams remained completely wild-type, with no transgenic protein disturbance in the intrauterine microenvironment. In the control group, wild-type male and female mice were mated, with neither offspring nor mothers carrying AD-related mutations, serving as a healthy baseline control. This design ensures that gestational changes more purely reflect pathological alterations during fetal development. Maternal urinary proteomic changes were dynamically monitored throughout pregnancy. This study aimed to characterize the ultra-early molecular response features of maternal mice during pregnancy under the background of offspring AD pathology and to screen for gestational AD-specific urinary differential proteins, with the goal of filling the research gap in non-invasive prenatal biomarkers for fetal AD.

## 2. Materials and Methods

### 2.1 Experimental Animals and Model Establishment

Specific pathogen-free (SPF) C57BL/6mice, including male 3xTg (APP/PS1/Tau) transgenic mice as well as wild-type male and female C57BL/6 mice, were purchased from Beijing HFK Bioscience Co., Ltd. Animal production license number: SCXK (Beijing) 2024-0003.All mice were housed in an SPF barrier facility under standardized environmental conditions: temperature maintained at 22–25 °C, relative humidity 50%–60%, and a 12-hour light/dark cycle. Standard rodent chow and sterile drinking water were provided ad libitum. All animal experiments were initiated after a one-week acclimatization period.

Animals were randomly divided into two groups, with 8 independent mating pairs set up for each group:Experimental group: 8-week-old male 3xTg-AD mice mated with wild-type female mice;Control group: 8-week-old wild-type male mice mated with wild-type female mice。Mating was conducted daily at 18:00 by housing male and female mice together at a male-to-female ratio of 2:1. Vaginal plugs in female mice were examined at 08:00 the following morning, and the day on which a vaginal plug was detected was recorded as gestational day 0.5 (GD 0.5).

### 2.2 Urine Sample Collection

Starting from 20:00 on the day of vaginal plug detection, pregnant mice were placed in metabolic cages overnight (from 20:00 to 8:00 the next morning) every 48 hours (on D1, D3, D5, D7, D9, D11, D13, D15, D17, and D19) for fresh urine collection. At least 500 μL of urine was collected from each mouse per time point. Immediately after collection, samples were centrifuged at 4°C and 3,000 r/min for 10 min to remove cell debris. The supernatant was aliquoted into sterile EP tubes and stored at −80°C, avoiding repeated freeze-thaw cycles.

### 2.3 Sample Processing

Protein extraction: Urine samples were thawed at 4°C and centrifuged at 12,000 ×g and 4°C for 10 min. The supernatant was mixed with pre-chilled absolute ethanol at a ratio of 1:3 (supernatant:ethanol) and precipitated at−20°C for 12 hours. After centrifugation at 12,000 × g and 4°C for 30 min, the supernatant was discarded. The pellet was dried and then dissolved in 500 μL of lysis buffer, sonicated for 3 min, and rotated at 4°C for 2 hours. Protein concentration was determined by the Bradford method.

Trypsin digestion: 100μg of protein was brought to 200μL with 25mM NH_4_HCO_3,_ and 4 μL of 1 M DTT (final concentration 20 mM) was added, followed by reduction in a water bath at 37°C for 1 h. After cooling, 10 μL of 1 M IAA (final concentration 50 mM) was added and the mixture was incubated in the dark at room temperature for 30 min for alkylation. Using a 10 kDa ultrafiltration tube, the membrane was washed twice with UA solution (8 M urea, 0.1 M Tris-HCl, pH 8.5). The alkylated protein sample was then loaded and centrifuged at 14,000 × g for 10 min, washed twice with UA solution, and rinsed three times with 200μL of NH_4_HCO_3_. Trypsin was added at an enzyme-to-protein ratio of 1:50 (w/w), and digestion was carried out overnight at 37°C. The next day, the filtrate (containing peptides) was collected by centrifugation at 4°C and 14,000×g for 30min. The peptides were desalted using an HLB solid-phase extraction cartridge, lyophilized, and stored at−20°C.

### 2.4 LC-MS/MS Mass Spectrometry Analysis

Lyophilized peptide pellets were reconstituted in 0.1% formic acid, and peptide concentrations were quantified using a BCA protein assay kit. Each sample was diluted to a concentration of 0.5 μg/μL. A 10 μL aliquot of each diluted sample was centrifuged at 12,000 g for 30 min at 4 °C, after which the supernatant was collected and spiked with 1 μL of IRT peptide standard. An equal volume (1 μL) was drawn from every sample and pooled to prepare a quality control (QC) mix for instrument monitoring prior to LC-MS/MS measurement.Label-free quantitative proteomic profiling was performed via liquid chromatography-tandem mass spectrometry (LC-MS/MS). Peptide separation was carried out on a nanoflow high-performance liquid chromatography (nano-HPLC) system, and mass spectra were acquired on a high-resolution tandem mass spectrometer to record full-scan MS and MS/MS data. All peptide samples were analyzed using an Orbitrap Exploris 480 mass spectrometer coupled to an UltiMate 3000 nano-LC system. The separation system was equipped with an integrated monolithic C18 capillary column (75 μm inner diameter, 50 cm length), and the column temperature was maintained at a constant 60 °C throughout the entire run.Peptides were separated with a segmented linear mobile phase gradient at a fixed flow rate of 1.5 μL/min over a total runtime of 25 min. Mobile phase B consisted of acetonitrile acidified with 0.1% (v/v) formic acid. The gradient program was set as follows: mobile phase B was increased from 5% to 20% within 15.5 min, ramped from 20% to 30% over the subsequent 5 min, elevated from 30% to 50% in 1 min, and rapidly raised to 90% within 0.1 min. The 90% B condition was held for 1.3 min, followed by a quick drop back to 5% B in 0.1 min, with a final 2 min re-equilibration at 5% B.The mass spectrometer was operated in positive electrospray ionization mode with data-independent acquisition (DIA) parameters configured as described below. For full MS survey scans: resolution = 120,000, automatic gain control (AGC) target = 300%, maximum ion injection time = 50 ms, mass scan range = 350–1200 m/z. For DIA MS/MS scans: resolution = 30,000, AGC target = 200%, maximum ion injection time = 50 ms. Higher-energy collisional dissociation (HCD) was applied with a normalized collision energy of 30%.

### 2.5 Data Search

The raw mass spectrometry files were searched using Spectronaut software (v19.0, Biognosys) against the Swiss-Prot mouse database. The enzyme specificity was set to Trypsin/P, allowing up to 2 missed cleavages, with peptide lengths ranging from 7 to 52 amino acids. Carbamidomethylation of cysteine was set as a fixed modification, while N-terminal acetylation and methionine oxidation were set as variable modifications, with a maximum of 5 variable modifications per peptide. The false discovery rate (FDR) at both the peptide and protein levels was set to ≤ 1%. Peptide abundance was calculated by summing the fragment ion peak areas in MS² spectra, and protein abundance was calculated by summing the abundances of its constituent peptides.

### 2.6 Data Analysis

The search results were normalized and missing values were imputed. Fold change (FC) was calculated using the group means to screen for differentially expressed proteins. The criteria for differentially expressed proteins were set as fold change (FC) ≥ 1.5 or ≤ 0.67, with a two-tailed unpaired t-test P-value < 0.05. For proteins with no quantitative signal in all samples of the control group, the fold change was recorded as ∞ (upregulated) or 0 (downregulated), indicating that the protein was detected exclusively in the experimental group or exclusively in the control group, respectively. To control for false positives, sample label permutation tests (random shuffling of group labels) were performed simultaneously to calculate the mean number of differential proteins in random groupings and the probability of obtaining the observed differential set by chance.

### 2.7 Bioinformatics Analysis

Systematic bioinformatic analyses were performed on the screened differentially expressed proteins (DEPs). Gene Ontology (GO) functional enrichment analysis was conductedusingtheDAVID database (https://davidbioinformatics.nih.gov/summary.jsp) to characterize the functional distribution of DEPs. Literature mining was carried out via the PubMed database (https://pubmed.ncbi.nlm.nih.gov) to identify core DEPs closely associated with AD pathological processes.

## 3. Results

### 3.1 Model Construction and Sample Collection

No statistically significant differences were observed between the two groups in pregnancy rates, gestational duration, or litter sizes (P>0.05). No maternal miscarriage or death occurred during the experimental period. All sequential urine samples showed no visible contamination or protein degradation, and protein concentrations and purities met the standards for mass spectrometry analysis, ensuring a reliable and balanced baseline. The QC samples showed stable coefficients of variation in repeated measurements, demonstrating good reproducibility of the LC-MS platform. No high-abundance exogenous contaminating proteins were detected in the empty metabolic cage negative controls.

### 3.2 Global Changes in the Gestational Urinary Proteome

A total of 2,729 proteins were identified across all time points (FDR < 1.0%), of which 1,250 proteins were retained for subsequent analysis after quality control filtering. Throughout embryonic development, differentially expressed proteins were present between the experimental and control groups at all gestational time points, suggesting that AD-related pathological changes may precede individual developmental maturation. The number of differential proteins exhibited stage-specific patterns (Figure 1): in early gestation (D1–D5), significantly upregulated proteins predominated, with almost no downregulated proteins; in mid-gestation (D9–D13), the number of upregulated proteins declined while downregulated proteins emerged in large numbers; in late gestation (D15–D17), upregulated proteins resurged alongside a large number of downregulated proteins; at D19, the total number of differential proteins dropped to its lowest level across the entire gestation (5 proteins).

**Figure 1.**
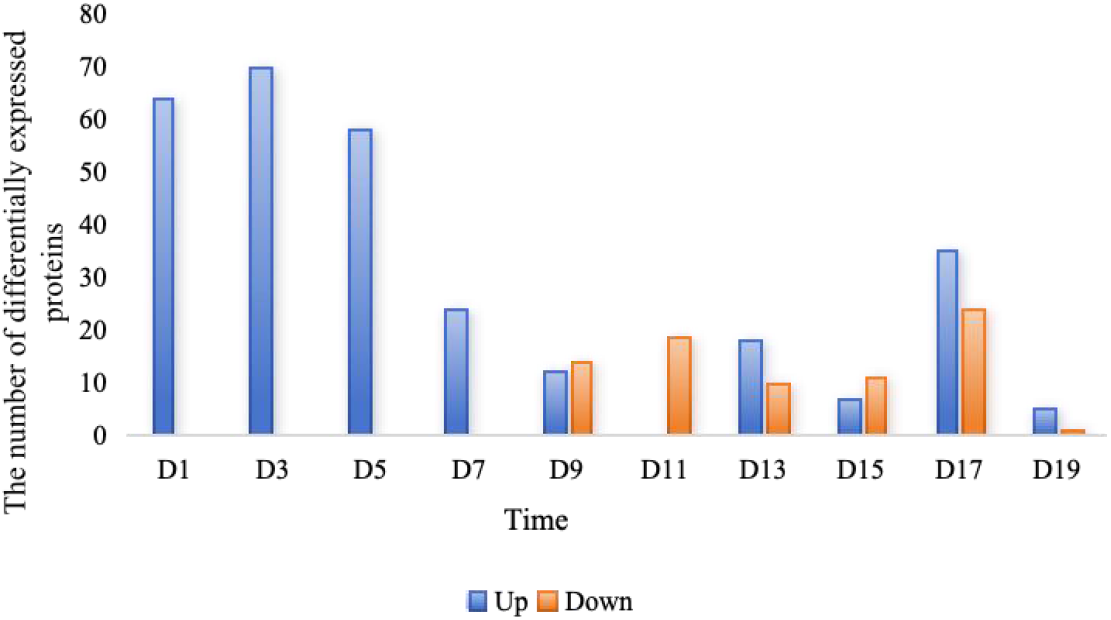
Distribution of differential protein)numbers at each time point (Blue: upregulated, FC ≥ 1.5, P < 0.05; Orange: downregulated, FC ≤ 0.67, P < 0.05)

### 3.3 Permutation Test Validation of Differential Protein Reliability

Using custom Python scripts, batch Welch’s t-tests (corrected for heteroscedasticity) were performed on the urinary protein quantitative data between the experimental and control groups at each gestational week, with differential screening thresholds of FC ≥ 1.5 or FC ≤ 0.67 and P < 0.05, without assuming homogeneity of variance, to avoid statistical bias caused by unequal intergroup dispersion in proteomic data^[26]^. To exclude false positives from random grouping, sample label permutation tests were performed simultaneously: group labels were randomly shuffled at each time point and the test was repeated, and the mean number of differential proteins in random groupings and the probability of obtaining the observed differential set by chance were calculated. The results are shown in Table 1.

**Table 1.** False positive assessment table for sample tests.

| Gestationnl<br>Time Point | Mean Proteins in Random<br>Groups | Total DEPS in Real<br>Experiment | Probability of Random<br>Generation |
| --- | --- | --- | --- |
| D1 | 2 | 42 | 0.048 |
| D3 | 2 | 38 | 0.053 |
| D5 | 1.4 | 26 | 0.054 |
| D7 | 0.8 | 14 | 0.057 |
| D9 | 0.87 | 16 | 0.054 |
| D11 | 0.7 | 14 | 0.050 |
| D13 | 0.8 | 13 | 0.062 |
| D15 | 0.5 | 12 | 0.042 |
| D17 | 0.95 | 13 | 0.073 |
| D19 | 0.208 | 5 | 0.042 |

The random permutation test results showed that across all gestational time points, the mean number of differential proteins detected in random groupings was only 0.21–2.00, far lower than the total number of differential proteins obtained from real experimental versus control comparisons(5–42). The probability of randomly generating the corresponding differential protein set at each time point was below 0.075 (with most below 0.06), indicating that the differential proteins detected in this study represent genuine biological signals rather than random statistical noise.

### 3.4 Functional Analysis of Differential Proteins

Differential proteins at each time point were subjected to protein functional analysis and biological enrichment using the DAVID database and PubMed.

Notably, at D1, 64 differential proteins were identified in the experimental group (Table 2), suggesting that the presence of paternal transgenes produces detectable molecular effects in the mother as early as the zygote formation and pre-implantation stage, and that this response can be directly reflected by urinary proteomic profiling. Among the 64 identified differential proteins, PubMed literature retrieval revealed that 23 were associated with neural pathways. Among these, the upregulation of Ras-related protein Rab-5A was particularly prominent. Rab-5A is a key member of the small GTPase Rab family and a well-established master regulator of the early endocytic pathway, primarily involved in the fusion of endocytic vesicles with early endosomes^[27]^. Previous reports indicate that RAB5A dysfunction also occurs in AD and other diseases. Its overactivation is thought to accelerate the production of amyloid-beta and may induce neuronal apoptosis^[28]^. Thus, the aberrant elevation of Rab5A strongly suggests hyperactivation of the endocytic pathway in the AD model, a change that can be captured by the urinary proteome.NAGPA, also known as mannose-6-phosphate exposure enzyme, is a key enzyme in the lysosomal enzyme sorting pathway^[29]^, and its expression level was significantly upregulated in the disease model. Currently, mainstream gene function databases (such as NCBI Gene and UniProt) and review articles have not reported a direct causal relationship between NAGPA and Alzheimer’s disease (AD). Given the widespread lysosomal/autophagy pathway dysfunction in AD, this change in NAGPA may suggest its involvement in the regulation of lysosomal homeostasis, potentially as a compensatory response to pathway stress. In addition, a transcriptomic study also observed NAGPA upregulation in the entorhinal cortex of AD patients, consistent with our proteomic findings^[30]^. This suggests that NAGPA may play a role in the pathophysiology of AD, although the specific mechanism warrants further investigation.Serine/threonine-protein kinase TAO1 (abbreviated as TAOK1) belongs to the MAP kinase kinase kinase (MAP3K) family^[31]^. Genome-wide association studies (GWAS) have shown that the TAOK1 gene is significantly associated with AD risk or family history and CSF Aβ1-42 levels^[32]^. Furthermore, the development of the first selective TAOK1 tool compound has further corroborated its status as a potential therapeutic target for AD^[33]^. Our results suggest that aberrant activation of TAOK1 may contribute to MAPK signaling dysregulation and cytoskeletal pathology in this model disease.Ubiquitin recognition factor in ER-associated degradation protein 1 (UFD1), as the ubiquitin-recognizing subunit of the VCP-NPLOC4-UFD1 complex, is responsible for retrotranslocating misfolded proteins from the ER lumen to the cytosol for targeted degradation in the ER-associated degradation (ERAD) pathway^[34]^. GWAS data suggest that UFD1 is associated with accelerated cognitive decline during the transition from mild cognitive impairment to Alzheimer’s disease^[35]^. Therefore, the significant upregulation of UFD1 in this model may reflect enhanced ER stress and compensatory cellular adaptation to proteotoxic stress.

**Table 2.** DEPS Identified at Gestational Day 1 in the Alzheimer’s Disease Model.

| ProteinAccessions | ProteinDescriptions | FC | Pvalue | Reference |
| --- | --- | --- | --- | --- |
| Q9CQD1 | Ras-related protein Rab-5A | ∞ | 4.42E-02 | {28, 29} |
| Q9LSF8 | Dimethylnonatriene synthase | ∞ | 4.45E-02 | |
| Q8BJ48 | N-acetylglucosamine-1-phosphodiester<br>alpha-N-acetylglucosaminidase | 127.28↑ | 2.12E-02 | {30} |
| Q5F2E8 | Serine/threonine-protein kinase TAO1 | 22.24↑ | 3.32E-02 | {31~33} |
| P70362 | Ubiquitin recognition factor in<br>ER-associated degradation protein 1 | 16.86↑ | 7.95E-03 | {34,35} |
| P01629 | Ig kappa chain V-II region 2S1.3 | 14.84↑ | 1.31E-02 |  |
| P11758 | Hemoglobin subunit beta | 14.66↑ | 4.81E-02 |  |
| Q9JIW9 | Ras-related protein Ral-B | 13.81↑ | 2.41E-02 |  |
| P21107 | Tropomyosin alpha-3 chain | 11.25↑ | 3.93E-02 |  |
| O89001 | Carboxypeptidase D | 10.89↑ | 2.03E-02 | {36} |
| Q9QYJ0 | DnaJ homolog subfamily A member 2 | 9.98↑ | 4.86E-02 | {37} |
| Q9QX15 | Calcium-activated chloride channel<br>regulator 3A-1 | 9.91↑ | 4.17E-02 |  |
| Q64105 | Sepiapterin reductase | 9.64↑ | 2.50E-03 | {38} |
| P62071 | Ras-related protein R-Ras2 | 9.62↑ | 4.41E-02 |  |
| Q8R121 | Protein Z-dependent protease inhibitor | 9.57↑ | 2.09E-02 |  |
| Q62426 | Cystatin-B | 9.34↑ | 7.92E-03 | {39} |
| P05063 | Fructose-bisphosphate aldolase C | 8.13↑ | 4.01E-02 | {40} |
| P10810 | Monocyte differentiation antigen CD14 | 8.07↑ | 2.96E-02 | {41} |
| P34902 | Cytokine receptor common subunit gamma | 7.96↑ | 4.61E-02 | {42} |
| P46467 | Vacuolar protein sorting-associated protein<br>4B | 7.67↑ | 1.24E-02 | {43} |
| Q9QZC7 | Pleckstrin homology domain-containing<br>family B member 2 | 7.46↑ | 4.96E-02 |  |
| Q80YC5 | Coagulation factor XII | 7.21↑ | 2.90E-02 |  |
| Q8VDL4 | ADP-dependent glucokinase | 6.58↑ | 4.74E-02 |  |
| P92549 | ATP synthase subunit alpha, mitochondrial | 6.46↑ | 1.82E-02 | {44} |
| P07758 | Alpha-1-antitrypsin 1-1 | 6.02↑ | 3.13E-02 |  |
| P60893;P60894 | Probable G-protein coupled receptor 85 | 5.65↑ | 2.01E-02 |  |
| Q9WTP6 | Adenylate kinase 2, mitochondrial | 5.63↑ | 4.00E-02 |  |
| Q8VEJ9 | Vacuolar protein sorting-associated protein<br>4A | 5.44↑ | 4.56E-02 |  |
| P17742 | Peptidyl-prolyl cis-trans isomerase A | 5.24↑ | 4.30E-02 | {45} |
| O35215 | D-dopachrome decarboxylase | 5.14↑ | 2.70E-02 |  |
| P48036 | Annexin A5 | 4.94↑ | 2.90E-02 | {46} |
| Q8BKX1 | BAR/IMD domain-containing adapter<br>protein 2 | 4.90↑ | 2.44E-02 |  |
| Q8R3P0 | Aspartoacylase | 4.69↑ | 2.64E-02 |  |
| P47757 | F-actin-capping protein subunit beta | 4.62↑ | 2.80E-02 |  |
| P45700 | Mannosyl-oligosaccharide<br>1,2-alpha-mannosidase IA | 4.51↑ | 3.60E-02 |  |
| P24638 | Lysosomal acid phosphatase | 4.50↑ | 3.17E-02 |  |
| Q8VHY0 | Chondroitin sulfate proteoglycan 4 | 4.15↑ | 1.16E-02 |  |
| Q9Z0P5 | Twinfilin-2 | 4.13↑ | 4.46E-02 |  |
| Q91VU0 | Protein FAM3C | 4.07↑ | 2.44E-02 | {<sup>47</sup>} |
| P01834 | Immunoglobulin kappa constant | 4.07↑ | 2.27E-02 |  |
| P08905 | Lysozyme C-2 | 3.96↑ | 3.55E-02 |  |
| P52760 | 2-iminobutanoate/2-iminopropanoate<br>deaminase | 3.90↑ | 4.48E-02 |  |
| P15948 | Kallikrein 1-related peptidase b22 | 3.88↑ | 1.65E-02 |  |
| Q93092 | Transaldolase | 3.79↑ | 4.81E-02 |  |
| P97328 | Ketohexokinase | 3.76↑ | 4.23E-02 |  |
| P49935 | Pro-cathepsin H | 3.71↑ | 4.64E-02 | {<sup>48</sup>} |
| O08992 | Syntenin-1 | 3.70↑ | 1.28E-02 | {<sup>49</sup>} |
| Q61411 | GTPase HRas | 3.68↑ | 3.00E-02 | {<sup>50</sup>} |
| E9Q3E1 | Aldehyde dehydrogenase family 3 member<br>B2 | 3.68↑ | 3.75E-02 |  |
| Q60866 | N-acetyltaurine hydrolase | 3.61↑ | 3.18E-02 |  |
| Q9DBF1 | Alpha-aminoadipic semialdehyde<br>dehydrogenase | 3.56↑ | 4.36E-02 |  |
| P97426 | Eosinophil cationic protein 1 | 3.34↑ | 4.86E-02 |  |
| Q91XE4 | N-acyl-aromatic-L-amino acid<br>amidohydrolase (carboxylate-forming) | 3.30↑ | 2.57E-02 |  |
| P18526 | Ig heavy chain V region 345 | 3.18↑ | 2.57E-02 |  |
| P46638;P62492 | Ras-related protein Rab-11B;Ras-related<br>protein Rab-11A | 3.17↑ | 4.83E-02 |  |
| Q8CG76 | Aflatoxin B1 aldehyde reductase member 2 | 3.16↑ | 4.47E-02 |  |
| P01868;P01869 | Ig gamma-1 chain C region secreted<br>form;Ig gamma-1 chain C region,<br>membrane-bound form | 3.11↑ | 4.66E-02 |  |
| P01887 | Beta-2-microglobulin | 2.87↑ | 3.41E-02 | {<sup>51</sup>} |
| P24452 | Macrophage-capping protein | 2.81↑ | 4.42E-02 |  |
| Q9WVK4 | EH domain-containing protein 1 | 2.73↑ | 3.74E-02 | {<sup>52</sup>} |
| Q9DBD0 | Inhibitor of carbonic anhydrase | 2.66↑ | 4.03E-02 |  |
| A2A8L5 | Receptor-type tyrosine-protein phosphatase<br>F | 2.62↑ | 2.06E-02 | {<sup>53</sup>} |
| P22953 | Heat shock 70 kDa protein 1 | 2.61↑ | 4.24E-02 | {<sup>54</sup>} |
| O09114 | Prostaglandin-H2 D-isomerase | 2.35↑ | 4.09E-02 |  |

On day 3, a total of 70 DEPS were identified, exhibiting dramatic expression changes that covered the core pathological mechanisms of AD, including vesicular/endocytic-multivesicular body transport systems, mitochondrial energy metabolism and oxidative stress defense, lysosomal proteolytic systems, calcium homeostasis and G-protein signaling regulation, as well as blood-brain barrier damage and neuroinflammation.Guanine nucleotide-binding protein G(i) subunit alpha-1 (GNAI1), an inhibitory G-protein alpha subunit and a nodal point in GPCR signal transduction, showed significantly altered phosphorylation levels in neuronal cell models treated with Aβ oligomers, suggesting its involvement in downstream Aβ signal amplification^[55]^. Plasma membrane calcium-transporting ATPase 1 (PMCA1), a key molecule for maintaining neuronal intracellular calcium homeostasis^[56]^, was detected exclusively under AD pathological conditions, indicating that cells were facing a severe calcium overload crisis and were forced to initiate de novo protein synthesis for extreme compensation. Previous studies have confirmed that both core pathological proteins of AD—β-amyloid (Aβ) and Tau—can directly inhibit PMCA enzymatic activity, and that methylene blue exerts neuroprotective effects by activating PMCA and blocking the inhibitory effects of Aβ and Tau. The extreme upregulation of PMCA1 reflects the cell’s final effort to counteract calcium homeostasis imbalance^[57]^.DnaJ homolog subfamily A member 2 (DNAJA2) belongs to the DnaJ/Hsp40 chaperone family and is an important molecule regulating Tau pathology^[58]^. Systematic screening studies have confirmed that overexpression of DNAJA2 significantly reduces Tau aggregation and intracellular levels^[59]^. Therefore, its massive induction under AD pathological conditions likely represents a core neuronal defense response against Tau toxicity and neurofibrillary tangle formation.NADH dehydrogenase [ubiquinone] 1 beta subcomplex subunit 6 (NDUFB6), as a component of mitochondrial respiratory chain complex I, has its encoding gene annotated in the Alzheimer’s disease pathway by authoritative databases such as KEGG, directly reflecting the cell’s attempt at drastic remodeling of the mitochondrial electron transport chain to cope with energy crisis under AD pathological stress^[60]^. The Rab scaffold protein IQGAP2, according to the specialized database ExonSkipAD, shows exon skipping events significantly associated with two key clinicopathological indicators of AD—Braak staging and the Clinical Dementia Rating (CDR)—suggesting that its alternative splicing regulation is closely related to AD progression, and the dramatic changes at the protein level warrant in-depth investigation^[61]^. The V-type ATPase, critical for lysosomal acidification, has its enzymatic activity inhibited by cleavage products of the amyloid precursor protein (APP), leading to lysosomal dysfunction. The dramatic upregulation of its subunits may represent a compensatory response to the lysosomal acidification crisis^[62,63]^, indicating that neurons initiate extreme compensatory repair programs in response to early Aβ toxicity. Although ATPase GET3 showed dramatic changes in the AD group, along with several other molecules with infinite upregulation and no quantitative signal in the control group—such as Lrpprc, US7Z, and PI5K7 proteins—the evidence for their involvement in AD is limited, and their expression may be related to stress-induced high expression under specific pathological conditions. Their specific pathological significance represents a direction worthy of future in-depth exploration.In the ultra-high upregulation group (FC≥100), Rab5A reached a 203.22-fold increase. As the earliest pathogenic endocytic regulatory molecule in AD, its aberrant overexpression drives early endosomal enlargement, abnormal APP processing, and massive generation of toxic Aβ^[55]^. Mitochondrial fatty acid oxidation protein ETFβ, phosphate transporter SLC25A3, and other molecules reflecting mitochondrial energy metabolism damage and exacerbated lipid peroxidation stress were also upregulated, with cells compensatorily activating mitophagy pathways^[64]^. In the mid-to-high upregulation range (20 ≤ FC < 100), multivesicular body ESCRT sorting proteins VPS4B, VTA1, and MVB12A, the key lysosomal protease Legumain, the anti-ferroptosis defense molecule GPX4, and the glucose transporter GLUT2 were enriched. These molecules synchronously mediate Aβ lysosomal clearance and pathological tau/Aβ exosome transmission across brain regions, representing core markers of early proteostatic imbalance in AD^[65,66].^ Mild and low-magnitude upregulated proteins included the blood-brain barrier damage marker Endoglin, the exosomal marker CD81, Eph/ephrin inflammatory regulatory pathways, and the mitochondrial regulatory protease CLPX, suggesting that as early as day 3 of the disease course, brain microvascular permeability elevation and glial pre-activation had already occurred^[65,67]^.Compared with the D1 differential protein profile, the upregulation magnitudes of all pathway molecules at D3 were significantly higher, and perturbation intensities of the three core pathological pathways—Rab-mediated endocytic dysfunction, mitochondrial oxidative stress, and lysosomal proteolysis—were amplified exponentially, completely replicating the molecular signature of ultra-early rapid progression in AD^[68,69]^. This confirms that high-fold differential proteins related to vesicular trafficking, mitochondrial metabolism, and lysosomal degradation can serve as time-specific non-invasive biomarkers for distinguishing different prodromal stages of the disease^[70]^. Detected exclusively in the AD group, the overall upregulation fold changes at D3 were substantially higher than those at D1, with further amplification of perturbations in mitochondrial, lysosomal, and small GTPase pathways, representing the continuous aggravation of early AD pathological damage and the initiation of stronger compensatory stress programs across all cellular systems.

**Table 3.**
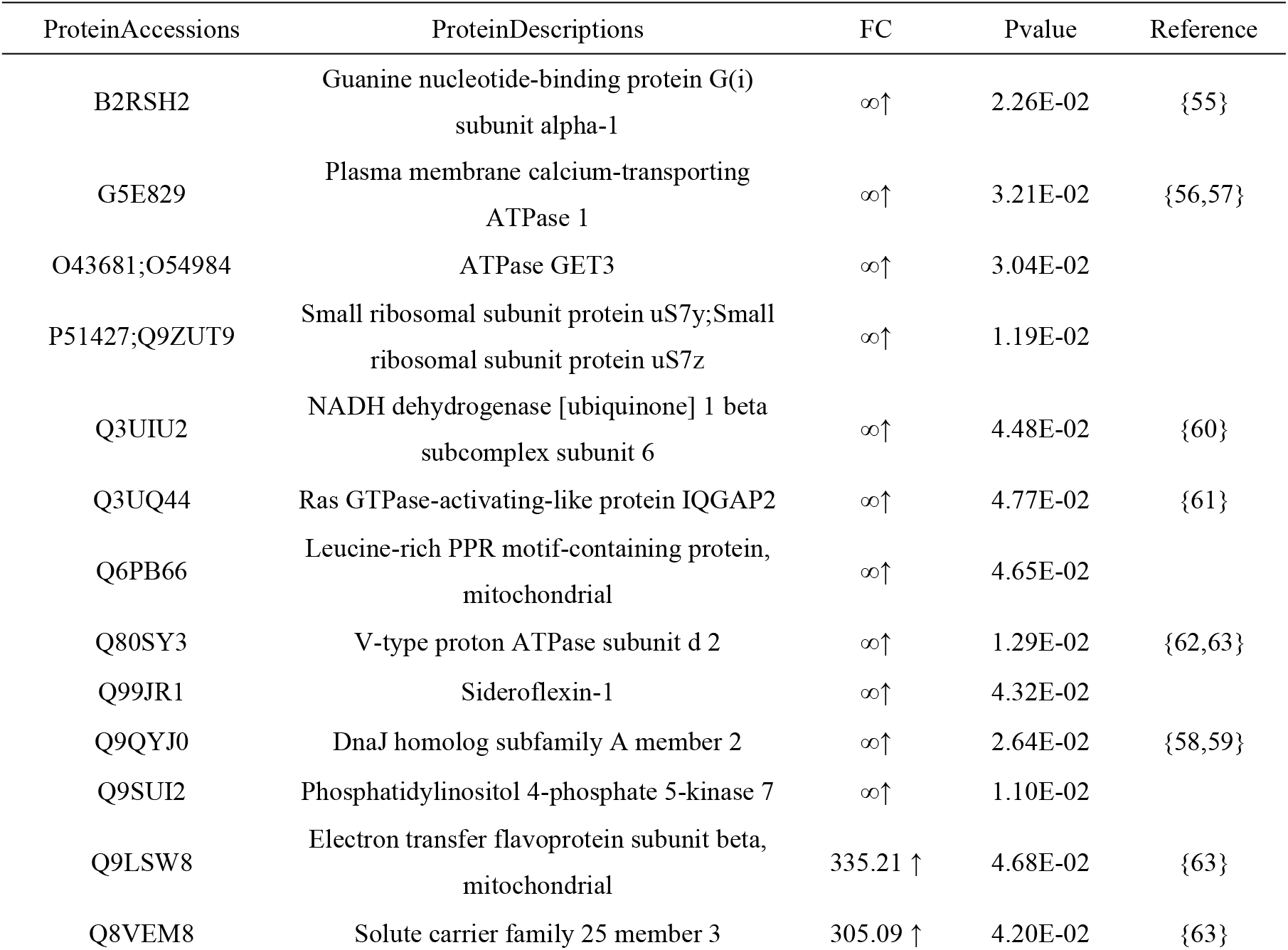
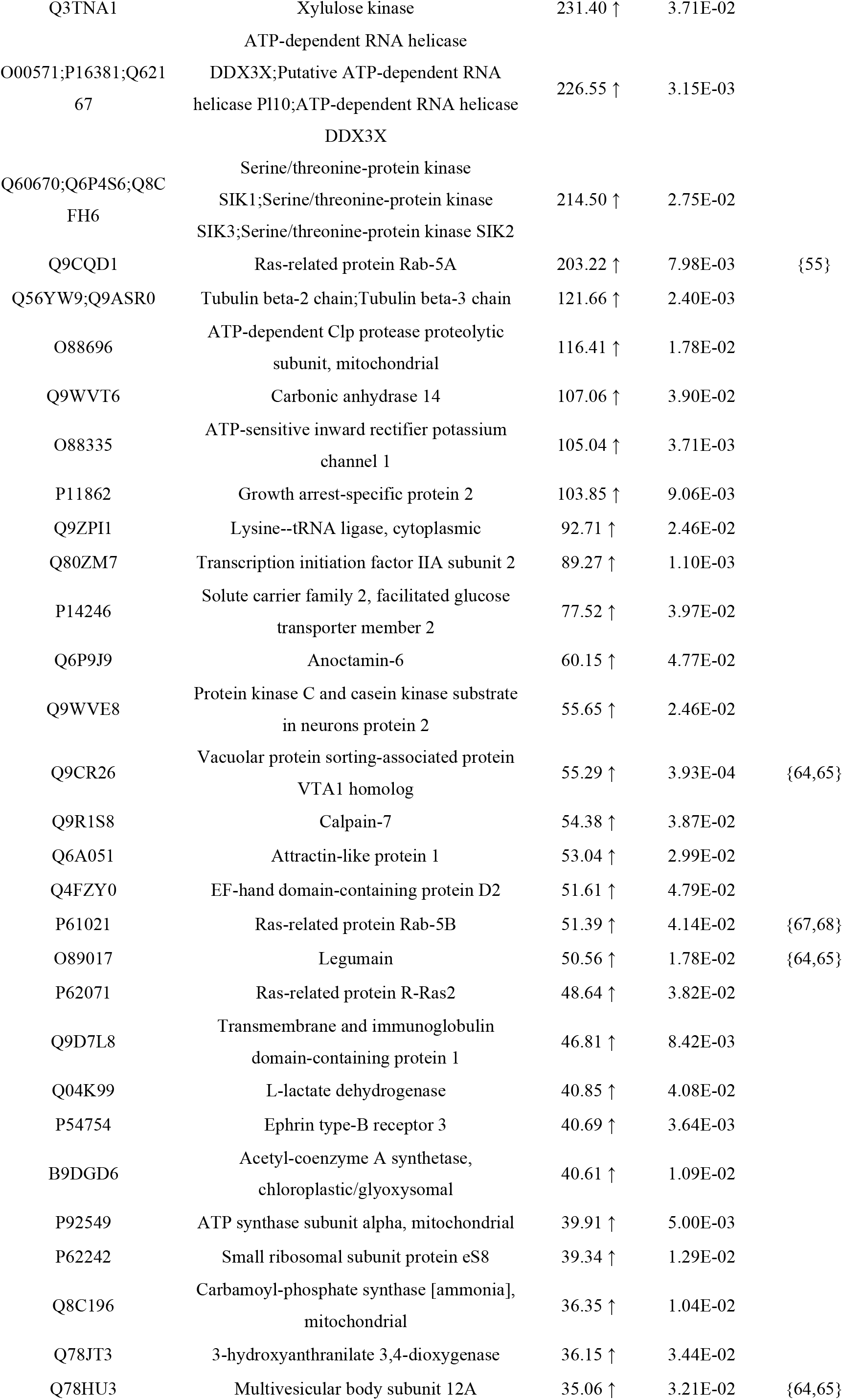
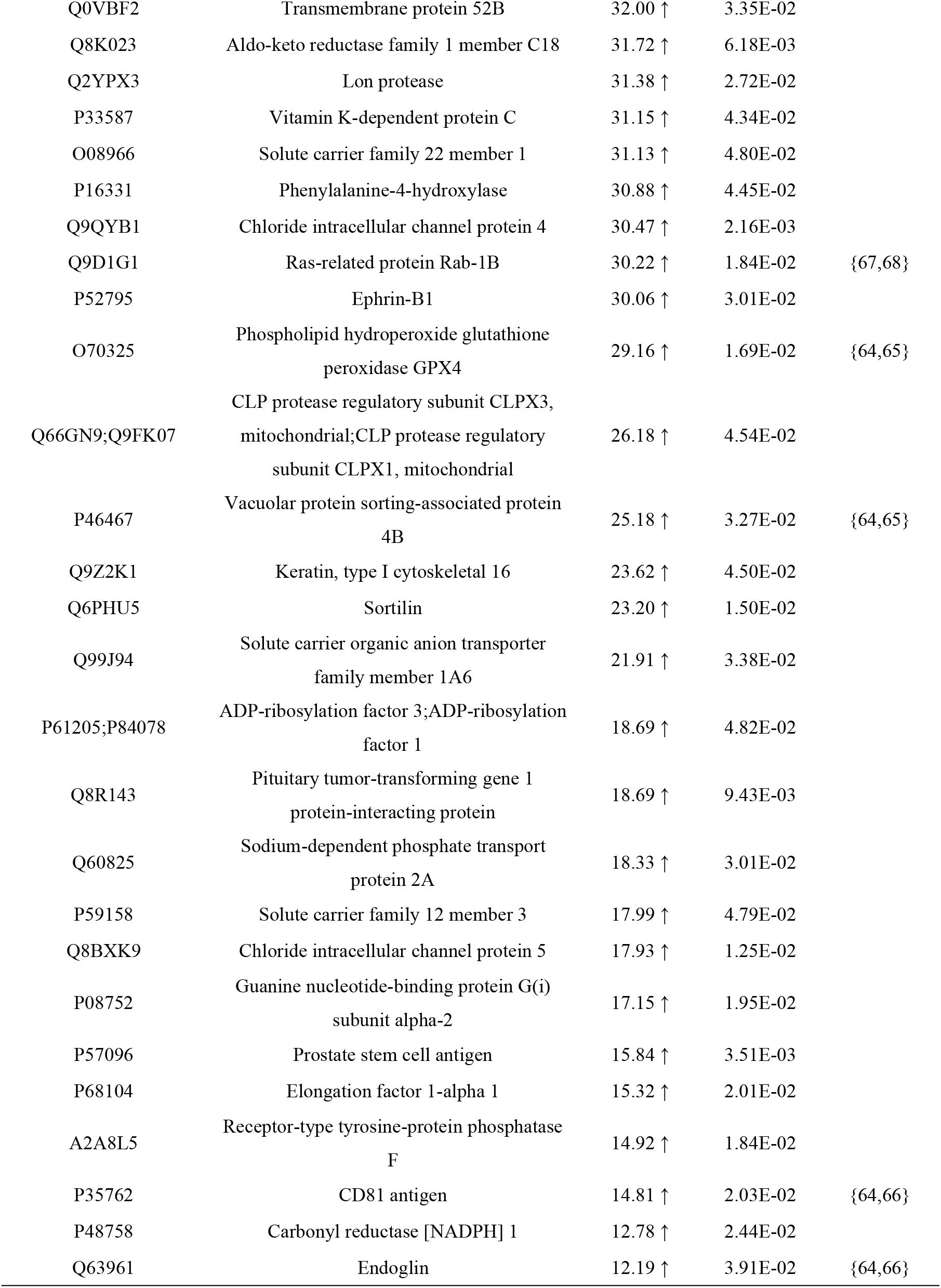
DEPS Identified at Gestational Day 3 in the Alzheimer’s Disease Model.

A total of 58 DEPs were identified at gestational day 5. These clustered proteins correspond to core AD pathological pathways, including extracellular matrix (ECM) remodeling, mitochondrial lipid metabolism, endocytic vesicle trafficking, neuroimmune inflammation, lysosomal proteolysis, and cytoskeletal synaptic damage. Among nine proteins exclusively detected in the experimental group, hyaluronidase 2 (HYAL2) and UDP-glucose dehydrogenase (UGDH) were simultaneously and extremely upregulated, strongly indicating severe disruption of hyaluronic acid (HA) metabolism and glycosaminoglycan (GAG) biosynthesis, which reflects systemic ECM remodeling under AD pathology. Degradation products of HA exhibit pro-neuroinflammatory activity, and HYAL2 itself is essential for maintaining blood-brain barrier integrity^[71]^. UGDH acts as the rate-limiting enzyme required for synthesizing precursors of HA and other GAGs such as chondroitin sulfate and heparan sulfate^[72]^. In AD, GAGs including heparan sulfate directly interact with Aβ and phosphorylated tau proteins, accelerating senile plaque and neurofibrillary tangle formation and modulating neuroinflammatory responses^[73]^. The synergistic extreme upregulation of HYAL2 and UGDH demonstrates systemic ECM remodeling in this AD mouse model, consistent with the well-documented abnormal deposition and degradation of ECM components in AD brains.TBC1 domain family member 10A (TBC1D10A), a negative Rab regulator, suppresses Rab5-mediated aberrant endocytosis to reduce APP cleavage and toxic Aβ generation^[74]^. Marked synchronous elevation of proteins involved in mitochondrial lipid anabolism and catabolism indicates progressive aggravation of Aβ-induced lipid peroxidation; cells activate fatty acid breakdown pathways to counteract neuronal ferroptosis and oxidative injury^[75]^. Increased copine-2 (CPNE2) reflects cellular stress responses to calcium overload. As Ca²⁺-dependent phospholipid-binding proteins, copines may exert vital regulatory roles in AD-associated disrupted calcium homeostasis^[76]^.Trichome birefringence-like protein 40 (TBL40,∞↑) has poorly characterized biological functions. Nevertheless, its extraordinarily high fold change implies its potential involvement in uncharacterized AD pathological cascades, requiring further validation in follow-up experiments.Phototropic-responsive NPH3 family protein NPY2 (178.68↑) shares homology with mammalian neurexophilin 3 (NXPH3), a neuron-specific secreted glycoprotein highly abundant in the brain. As a ligand of α-neurexin, NXPH3 participates in neuropeptide-like signal transduction and sustains synaptic architecture^[77]^.Recent genetic research has verified that the single-nucleotide polymorphism rs2234759 located in the promoter region of neuropeptide Y receptor Y2 gene (NPY2R) is strongly associated with AD; carriers of the G allele exhibit a 4.39-fold higher AD risk, and this variant may potentiate inhibitory control over presynaptic glutamate release^[78]^.At the mitochondrial level, extreme upregulation of 3-oxoacyl-[acyl-carrier-protein] synthase (OXSM) and 2-hydroxyacyl-CoA lyase 1 (HACL1) directly reveals severe dysregulation of mitochondrial fatty acid synthesis and α-oxidation pathways. OXSM mediates mitochondrial fatty acid biogenesis, while HACL1 serves as a key enzyme in phytanic acid α-oxidation; aberrant activity of both proteins is tightly linked to mitochondrial energy failure in AD^[79,80]^.Robust upregulation of cytoskeletal proteins (multiple keratins, actin) and desmosomal/hemidesmosomal junction proteins (plakophilin-1, desmoplakin, plectin), together with significant elevation of lysosomal markers LAMP1/LAMP2 and ESCRT complex subunit CHMP5, collectively demonstrate progressive systemic cellular structural collapse and decompensation of degradative clearance systems in the AD model at this gestational time point.Previous studies have validated that urinary proteomics recapitulate cytoskeletal pathological alterations characteristic of neurological disorders. For instance, AD7c-NTP, an AD-associated neurofilament protein, is markedly elevated in urine from AD patients and positively correlated with cognitive impairment severity^[81]^. Urinary exosome proteomic profiling of AD model mice also revealed prominent enrichment of proteins governing Aβ metabolism and clearance, matching the trends of altered endolysosomal and cytoskeletal proteins observed in the present study^[82]^. Urinary extracellular vesicle (EV) proteomes contain abundant endolysosomal proteins linked to AD, suggesting urinary EVs represent a non-invasive reservoir of biomarkers mirroring cerebral pathological lesions^[83]^. Our findings align with prior reports showing elevated LAMP2 in AD, accompanied by preferential routing of multivesicular bodies toward lysosomal degradation rather than secretion^[74,84]^. Differential expression of proteins such as GPRC5B has also been documented in urine samples from AD patients^[85]^.Collectively, these results confirm that urinary proteomics can capture multi-system pathological perturbations driven by AD, and further highlight ECM metabolism, calcium signaling and mitochondrial function as potentially critical driving pathways governing AD progression.

**Table 4.**
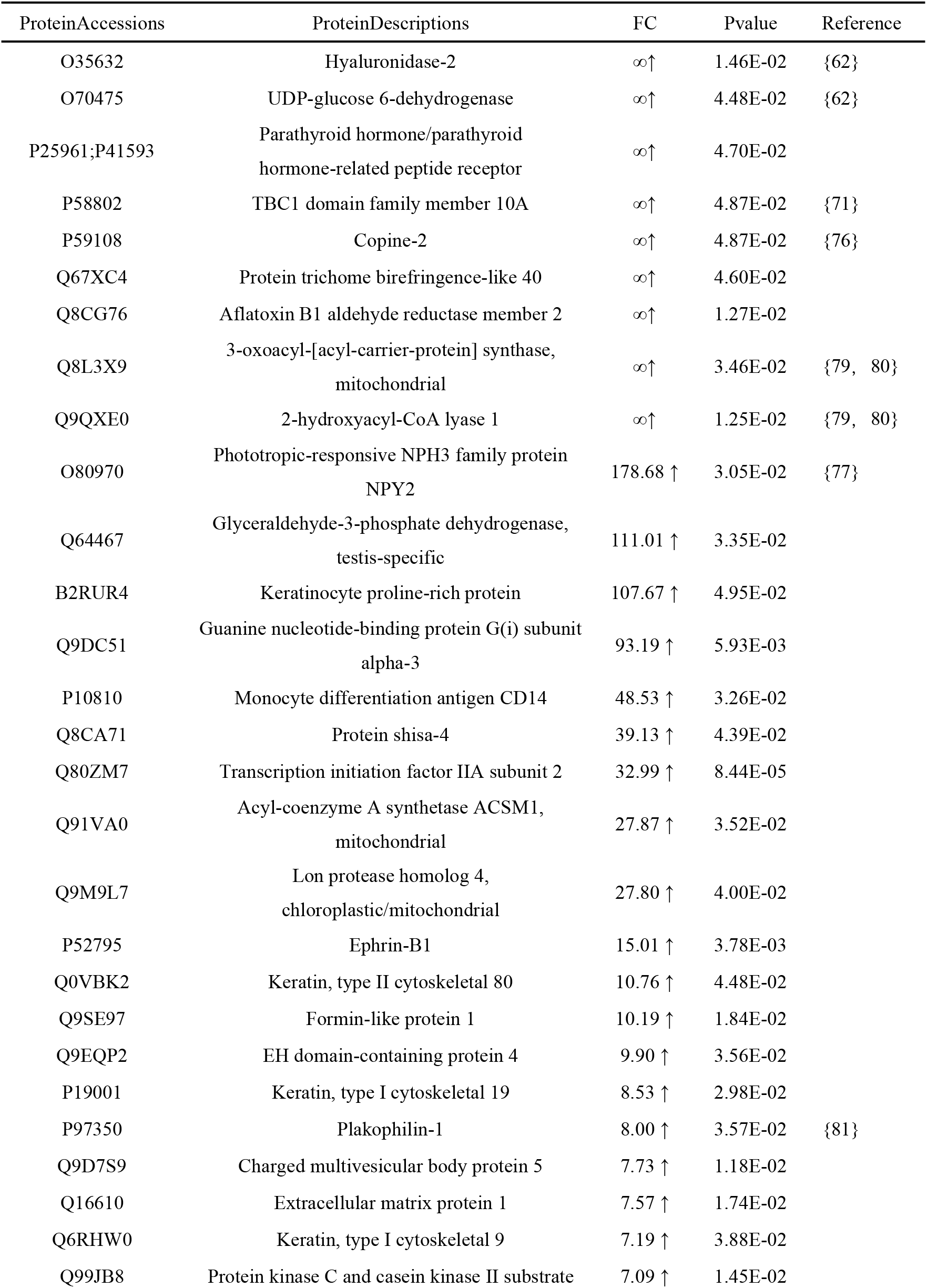
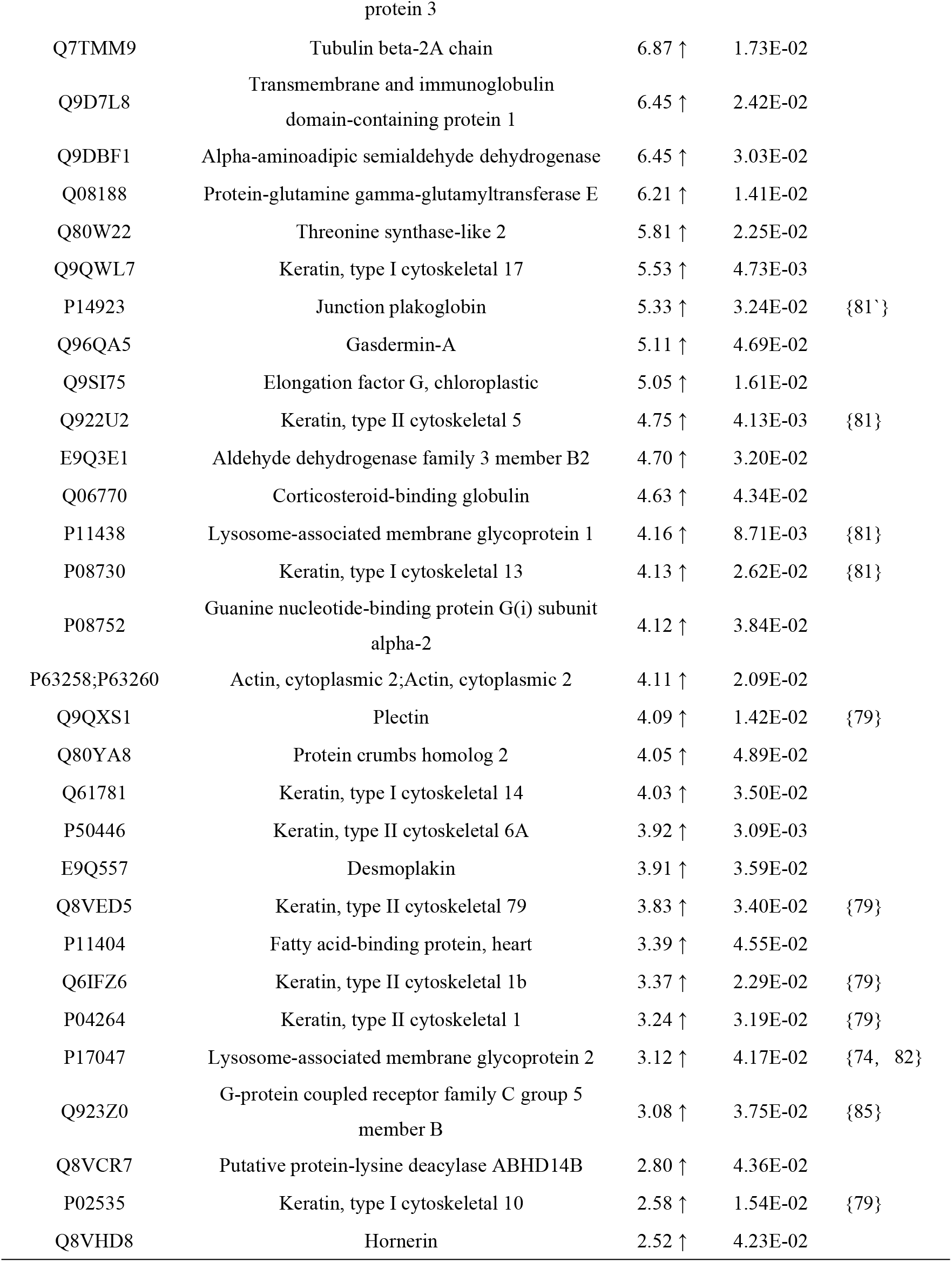
DEPS Identified at Gestational Day 5 in the Alzheimer’s Disease Model.

Urinary proteomic profiling at gestational day 7 revealed an evidently reduced number of DEPs compared with earlier time points (gestational days 1, 3 and 5), yet these DEPs were highly functionally concentrated.One of the most prominent findings at gestational day 7 was the marked upregulation of amyloid beta precursor-like protein 2 (APLP2, 36.00 ↑). As a vital member of the amyloid precursor protein (APP) family, APLP2 shares high structural homology and proteolytic processing patterns with APP: it is predominantly cleaved by β-secretase and γ-secretase to generate Aβ-like metabolites, and its aberrant metabolism may directly participate in AD-associated pathological cascades^[86]^. In addition, the extracellular domain of APLP2 has been validated as a master modulator of neuronal calcium homeostasis; simultaneous loss of APP and APLP2 impairs endoplasmic reticulum calcium store refilling and synaptic plasticity^[87]^. Existing evidence demonstrates that both APP and APLP2 are indispensable for neuromuscular transmission, spatial learning and synaptic plasticity, and their synergistic interplay is required to sustain intact neural function[88]. Dysregulated expression or dysfunction of APLP2 has been linked to AD pathogenesis, and its robust urinary upregulation may directly reflect severe perturbation of the central nervous system APP-family protein metabolic network.Another core observation was the significant elevation of ubiquitin-like protein NEDD8 (10.04↑). NEDD8 modulates the activity of cullin-RING E3 ubiquitin ligases via neddylation modification, thereby governing overall functionality of the ubiquitin-proteasome system. Immunohistochemical studies have confirmed that NEDD8 co-localizes with ubiquitin within neurofibrillary tangles and senile plaques in AD patients, indicating its direct involvement in the turnover of canonical pathological protein aggregates in AD. The proinflammatory cytokine IL-1β further drives nucleocytoplasmic translocation of NEDD8 and facilitates parkin-mediated ubiquitination,mechanism validated in the hippocampus of AD patients^[89,90]^.ADGRF5 belongs to the adhesion G protein-coupled receptor (aGPCR) family, whose members primarily mediate cell-cell and cell-extracellular matrix interactions. Dysfunctional GPCR signaling is tightly implicated in the initiation and progression of central nervous system disorders, and its sharp upregulation likely reflects progressive deterioration of intercellular communication and synaptic maintenance during AD pathogenesis^[91].^Carboxypeptidase E (CPE, 4.74↑) acts as a key carboxypeptidase governing neuroendocrine peptide maturation, responsible for removing C-terminal basic residues from precursor proteins inside secretory vesicles^[92]^. Nicotinamide phosphoribosyltransferase (NAMPT, 3.25-fold↑) is the rate-limiting enzyme of the NAD⁺ salvage biosynthesis pathway. Sustained NAD⁺ homeostasis is critical for neuronal wellness, and enhanced NAMPT activity elevates intracellular NAD⁺ concentrations to exert neuroprotective effects^[93]^. The coordinated upregulation of CPE and NAMPT may represent a dual compensatory strategy adopted by neurons under AD pathological stress: neurons attempt to reinforce neuropeptide signaling to counter synaptic dysfunction, while simultaneously boosting NAD⁺ synthesis to resolve energy deficit and oxidative stress.Collectively, urinary proteomic data obtained at gestational day 7 are highly focused on core pathogenic mechanisms underlying Alzheimer’s disease.

**Table 5.**
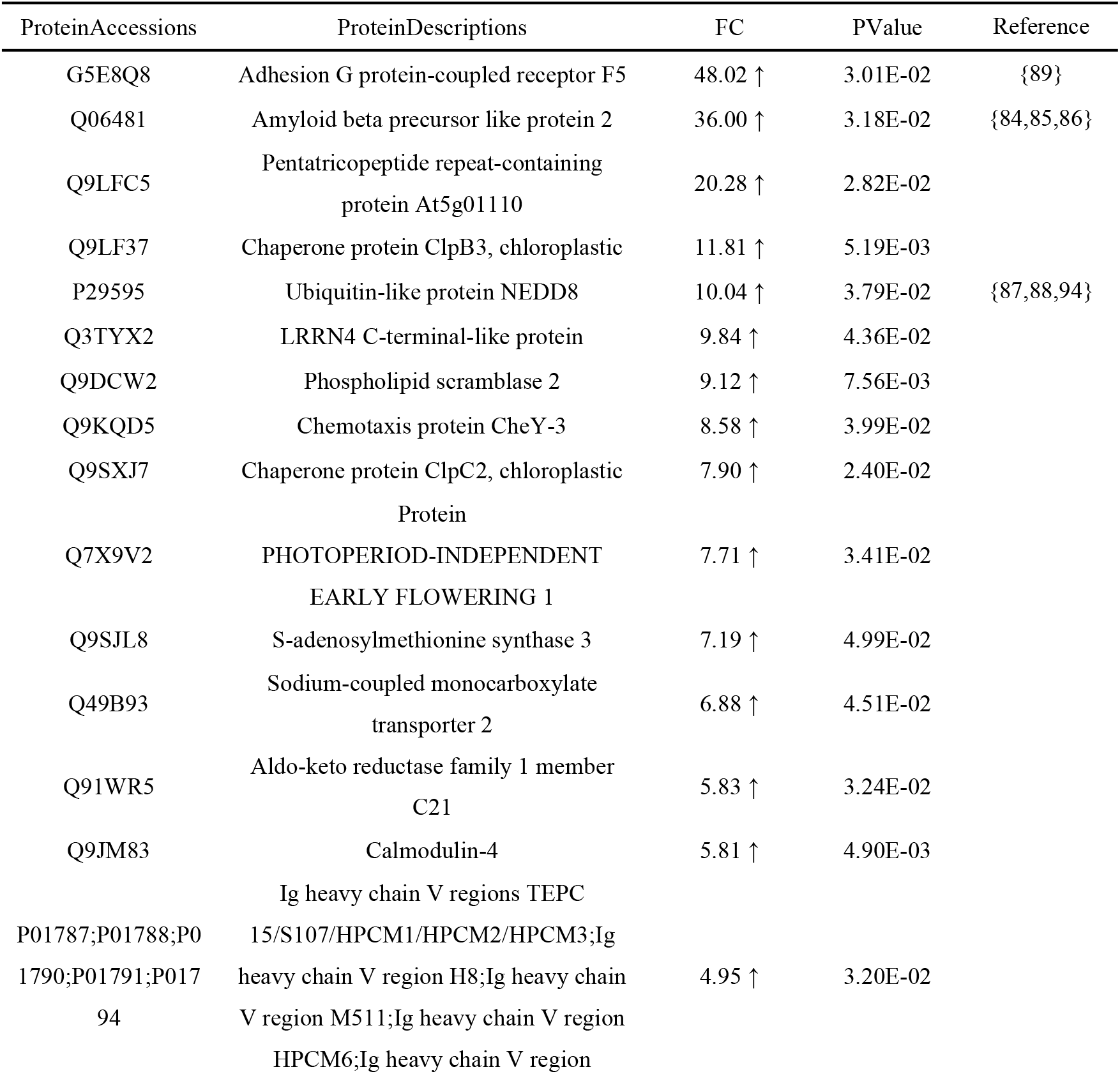
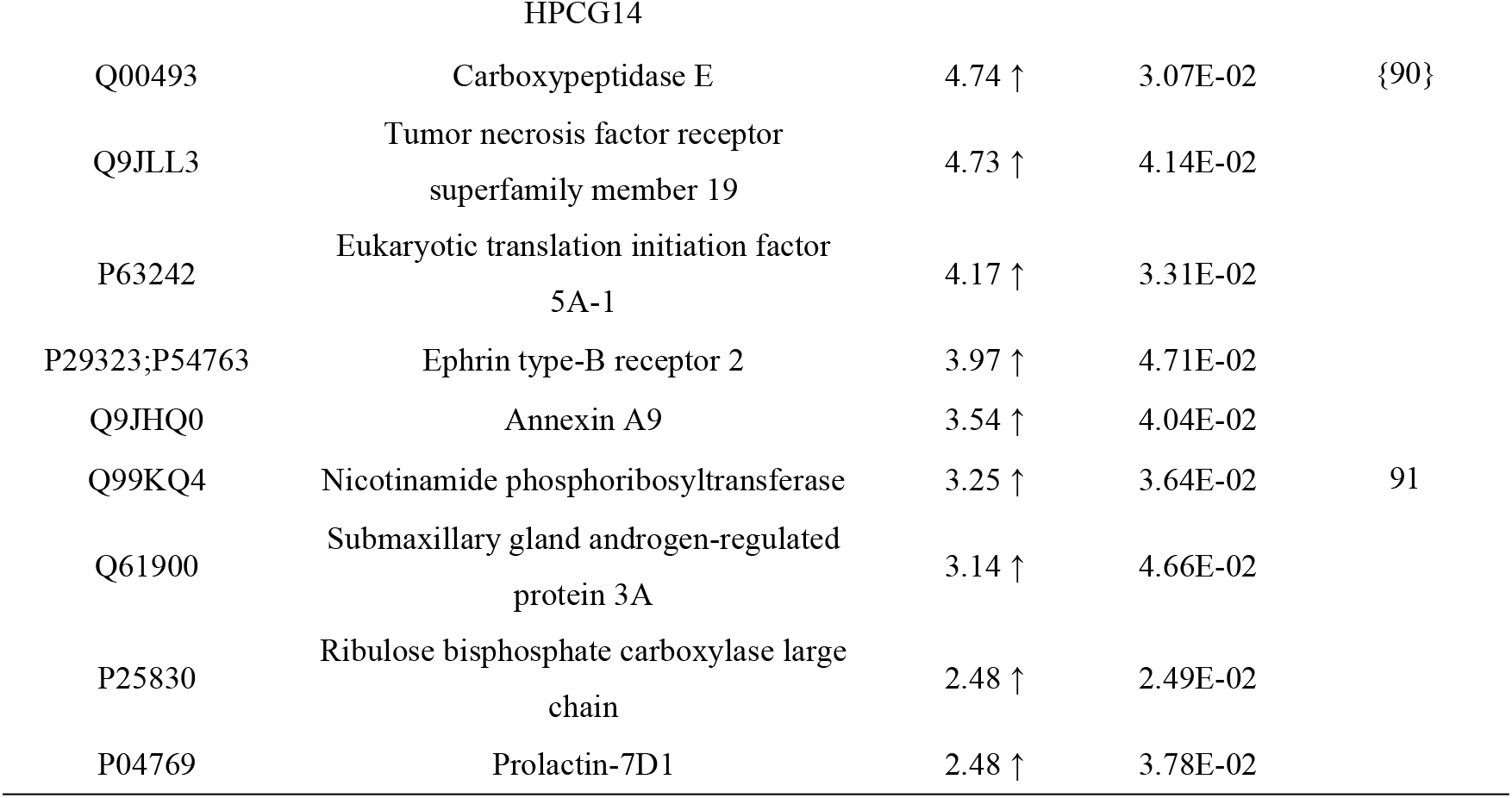
DEPS Identified at Gestational Day 7 in the Alzheimer’s Disease Model.

On day 9, the urinary proteome exhibited a pattern distinctly different from earlier stages—the number of differential proteins continued to decrease, but a large number of significantly downregulated proteins appeared for the first time. The most dramatic change was observed for Phototropic-responsive NPH3 family protein NPY2 (∞↑). The mammalian homolog of this protein is Neurexophilin 3 (NXPH3), a neuron-secreted glycoprotein highly expressed in the brain that participates in neuropeptide-like signal transduction and synaptic structure maintenance as a ligand for alpha-neurexin^[77]^. Notably, NPY2 had already shown a 178.68-fold upregulation at day 5 and further increased to infinite upregulation by day 9, suggesting sustained intensification of the activation of this synapse-related signaling pathway ^[78]^.Tropomyosin alpha-3 chain (TPM3, 10.61↑), an actin-binding protein involved in stabilizing the cytoskeleton and synapse formation, may reflect compensatory responses to maintain cytoskeletal integrity in AD^[94,95]^.Lysosomal alpha-mannosidase (MAN2B1, 8.01↑), a key enzyme in lysosomal degradation of mannose-containing glycoproteins, suggests compensatory activation of the lysosomal system, consistent with reports of enhanced autophagy-lysosomal pathways for clearing aberrant protein aggregates in AD^[96]^. The coordinated upregulation of Histone H2A (3.58↑) and Actin-like protein 6A (ACTL6A, 2.96↑) suggests activation of chromatin remodeling complexes^[97]^. The upregulation of Sarcoplasmic/endoplasmic reticulum calcium ATPase 2 (SERCA2,2.66↑) is consistent with calcium homeostasis imbalance theories in AD and may represent a compensatory neuronal response to cytosolic calcium overload^[98]^. The elevation of Myelin basic protein (MBP, 2.00↑) may reflect white matter damage and myelin repair responses in AD^[99]^.However, the most critical pathological signals at day 9 came from the sharp downregulation of numerous core functional proteins. The severe downregulation of Tubulin alpha-4A chain (TUBA4A, 0.11↓) and F-actin-capping protein subunit alpha-2 (CAPZA2, 0.17↓) marked a comprehensive collapse of microtubule and actin networks—TUBA4A mutations have been linked to neurodegenerative diseases, and microtubule instability is an important cause of axonal transport deficits in AD^[100]^. The sharp decline of Thrombospondin-1 (THBS1, 0.05↓) and Collagen alpha-2(IV) chain (COL4A2, 0.46↓) indicates that the extracellular matrix was undergoing severe catabolism. THBS1 is aberrantly expressed in AD brains and affects synaptic maintenance; multi-tissue methylation analysis has identified THBS1 as one of the key AD-associated genes^[101]^. COL4A2, as a major component of basement membranes, its downregulation may weaken the natural inhibition of Aβ aggregation, and the AD Knowledge Portal has confirmed its differential expression in AD brains^[102]^. The complete loss of MUC6 (0.00↓) has profound pathological implications, as its genetic polymorphisms have been significantly associated with tau pathology severity in AD^[103]^. ADGRL1 (0.28↓), a presynaptic adhesion molecule, has its downregulation closely linked to synaptic function loss, and the AD Knowledge Portal clearly shows its significant differential expression at both RNA and protein levels in AD brains^[104,105]^. CADM1 (0.37↓), also an adhesion molecule, has been confirmed to cause synaptic functional defects upon its loss of function^[106]^. ACOX1 (0.26↓), the rate-limiting enzyme of peroxisomal β-oxidation, showed downregulation in the AD model accompanied by compensatory increases in peroxisome density and enzyme expression, suggesting exacerbation of lipid metabolism disturbances^[107]^.Collectively, these findings suggest that the urinary proteome can not only reflect the dynamic progression of AD-related systemic pathology but also capture the “tipping point” signal marking decompensation as early as day 9.

**Table 6.**
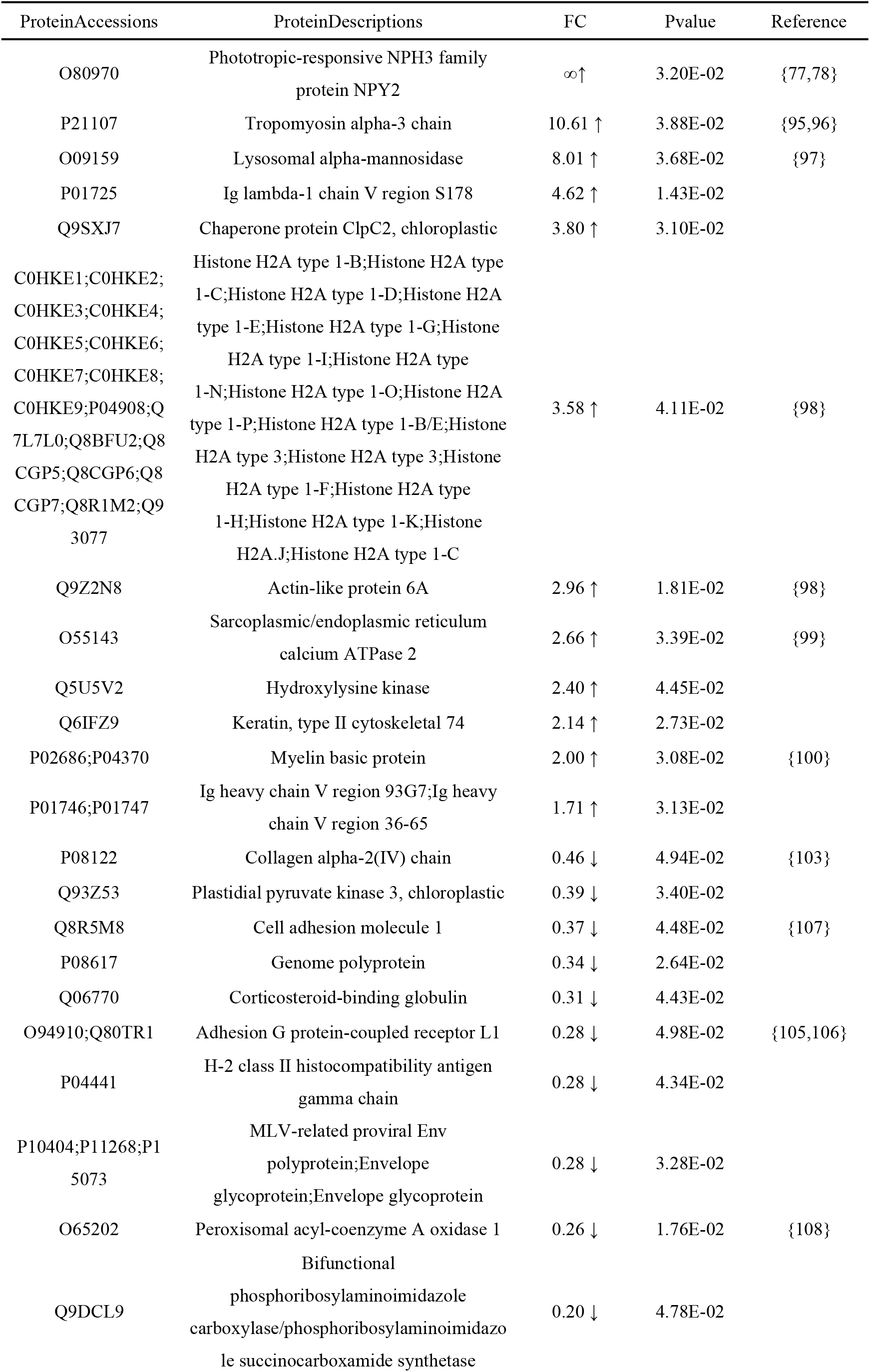
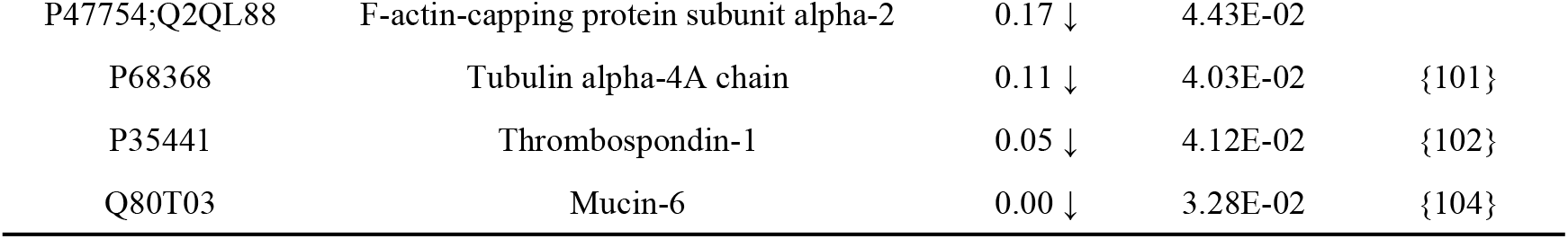
DEPS Identified at Gestational Day 9 in the Alzheimer’s Disease Model.

On day 11, the urinary proteome exhibited a pattern distinctly different from earlier stages: all differential proteins showed a downward trend, with extremely pronounced downregulation magnitudes, and more than half of the proteins had FC values below 0.3. Among these, the complete loss of Ezrin (0.00↓) was a notable signal. Ezrin is a core member of the ERM (ezrin-radixin-moesin) protein family, responsible for linking the plasma membrane to the actin cytoskeleton and regulating cell morphology, adhesion, and migration. ERM family proteins have been shown to be closely associated with activated microglia surrounding amyloid plaques in AD mouse models, and their expression changes directly participate in neuroinflammatory regulation^[108]^. The extreme downregulation of Desmoplakin (0.03↓) marked the disassembly of desmosomal structures. Recent studies have also shown that Dsp is specifically expressed in the hippocampal dentate gyrus and participates in maintaining neuronal activity and adult neurogenesis^[109]^. In this study, the very low level of Desmoplakin in urine (0.03↓) may reflect the terminal depletion of desmosomal proteins from the peripheral circulation in AD.Rho GDP-dissociation inhibitor 2 (ARHGDIB, 0.34↓), a key negative regulator of Rho GTPases, maintains Rho proteins in their inactive state and regulates cell migration, adhesion, and cytoskeletal organization. Its downregulation leads to aberrant activation of Rho signaling pathways, further exacerbating cytoskeletal and synaptic structural disruption, and it has been listed as an AD nominated target by Agora^[110]^. Glia maturation factor beta (GMFB, 0.44↓) is a brain-specific protein involved in the regulation of neuronal and glial maturation. Multiple studies have confirmed that GMFB is significantly elevated in all affected brain regions of AD patients and is considered an important participant in AD pathogenesis^[111].^Vacuolar protein sorting-associated protein 4A (VPS4A, 0.44↓), a core ATPase of the ESCRT pathway, has been identified as a differentially expressed gene related to vesicular transport and autophagy in an AD tau seeding model^[112]^.The collapse-level downregulation of Low-density lipoprotein receptor(LDLR, 0.06↓) has profound pathological significance—LDLR participates in cerebral Aβ clearance through binding to APOE, and its overexpression significantly reduces Aβ plaque deposition^[113,114]^. The extreme downregulation of Angiotensin-converting enzyme (ACE, 0.05↓)is directly associated with increased AD risk; multi-omics analysis has confirmed that genetically reduced ACE expression promotes AD through exacerbated tau phosphorylation, and ACE differential expression was also detected in urinary exosome proteomics of 5XFAD mice^[19,115]^. The severe downregulation of Fatty acid-binding protein (FABP1, 0.06↓) and Acetyl-coenzyme A synthetase (ACSS2, 0.20↓) reflects global suppression of lipid metabolism and energy production—ACSS2 has been identified as a differentially expressed gene related to metabolic reprogramming in AD models^[112]^.

**Table 7.** DEPS Identified at Gestational Day 11 in the Alzheimer’s Disease Model.

| ProteinAccessions | ProteinDescriptions | FC | Pvalue | Reference |
| --- | --- | --- | --- | --- |
| E9Q612 | Receptor-type tyrosine-protein phosphatase O | 0.57 ↓ | 3.85E-02 |  |
| Q9CQI3 | Glia maturation factor beta | 0.44 ↓ | 4.12E-02 | {112} |
| P97369 | Neutrophil cytosol factor 4 | 0.44 ↓ | 1.57E-02 |  |
| Q8VEJ9 | Vacuolar protein sorting-associated protein 4A | 0.44 ↓ | 3.06E-02 | {113} |
| Q6IMF0 | Keratin, type II cuticular 87 | 0.37 ↓ | 1.17E-02 |  |
| Q61599 | Rho GDP-dissociation inhibitor 2 | 0.34 ↓ | 4.16E-02 | {111} |
| Q99J77 | N-acetylneuraminate-9-phosphate synthase | 0.30 ↓ | 2.95E-02 |  |
| Q91V64 | Isochorismatase domain-containing protein 1 | 0.29 ↓ | 3.49E-02 |  |
| P63323 | Small ribosomal subunit protein eS12 | 0.26 ↓ | 4.37E-02 |  |
| Q93Z53 | Plastidial pyruvate kinase 3, chloroplastic | 0.22 ↓ | 1.71E-02 |  |
| Q99L13 | 3-hydroxyisobutyrate dehydrogenase, mitochondrial | 0.21 ↓ | 2.32E-02 |  |
| B9DGD6 | Acetyl-coenzyme A synthetase, chloroplastic/glyoxysomal | 0.20 ↓ | 3.70E-03 | {113} |
| Q8CIH5 | 1-phosphatidylinositol 4,5-bisphosphate phosphodiesterase gamma-2 | 0.11 ↓ | 1.39E-02 |  |
| Q810J1 | Pregnancy-specific glycoprotein 22 | 0.08 ↓ | 1.05E-02 |  |
| P35951 | Low-density lipoprotein receptor | 0.06 ↓ | 3.21E-02 | {114,115} |
| P12710 | Fatty acid-binding protein, liver | 0.06 ↓ | 3.93E-02 | {113} |
| P09470 | Angiotensin-converting enzyme | 0.05 ↓ | 1.78E-02 | {19,116} |
| E9Q557 | Desmoplakin | 0.03 ↓ | 2.61E-02 | {110} |
| P26040 | Ezrin | 0.00 ↓ | 6.31E-03 | {109} |

On day 13, as a mid-to-late gestational time point, the urinary proteome exhibited a unique pattern of “extreme activation coexisting with systemic decline.” On one hand, multiple proteins showed infinite upregulation or hundreds-fold increases, reflecting that endocytic-lysosomal clearance and calcium signaling pathways were in a state of overload stress; on the other hand, numerous metabolic enzymes and structural proteins showed severe downregulation or even complete disappearance.Among the upregulated proteins, the infinite upregulation of ADAM15 (∞↑)was the most prominent.ADAM15 belongs to the disintegrin and metalloproteinase family and is recorded in the ExonSkipAD database as a differentially expressed and exon-skipping gene associated with AD, suggesting transcript-level abnormalities in AD brain tissue^[116]^. Mendelian randomization studies of the CSF proteome have further suggested a potential causal role for ADAM15 in neurodegenerative diseases, and GWAS studies have identified it as an AD-associated candidate gene^[117,118]^. Anoctamin-6 (ANO6, 121.80↑) regulates ADAM10 activity through the induction of phosphatidylserine exposure; ADAM10, as an α-secretase, participates in non-amyloidogenic APP processing, and its dysregulation directly contributes to AD pathogenesis^[119]^. Sorcin (24.91↑) is highly expressed in the human brain, is significantly elevated in AD models, and interacts with the endoplasmic reticulum calcium channels RyR2/RyR3. Its upregulation may reflect severe intracellular calcium stress in the context of increased gestational calcium demand^[120]^. The elevation of PLD3 (4.05↑) directly points to core AD pathology—its genetic polymorphisms are associated with late-onset AD risk, and AD-associated variants can lead to aberrant APP processing and elevated Aβ levels^[121]^. CHMP5 (8.13↑), as a core component of ESCRT-III, has evidence of RNA differential expression and brain eQTL in AD brain tissue^[122]^.Translational regulator CsrA (∞↑) is a global carbon storage regulatory protein in prokaryotes. The direct association between CsrA and neurological diseases is currently very limited, and no clear evidence has been identified in published literature. Chaperonin GroEL 2 (∞↑), also known as heat shock protein 60, is closely associated with neurodegenerative diseases and can exert neuroprotective effects in AD and Huntington’s disease by inhibiting Aβ42 aggregation and reducing polyglutamine toxicity^[123]^.The complete disappearance of Pancreatic alpha-amylase 2a5(0.00↓), along with the severe downregulation of Sulfotransferase 1C2 (SULT1C2, 0.05↓) and Rab GDP dissociation inhibitor beta (GDI2, 0.26↓), may represent the collapse of metabolic enzyme systems and vesicular transport regulatory networks. GDI2 knockout has been shown to alleviate neurodegeneration and memory loss in 5xFAD mice^[124]^. Pancreatic alpha-amylase 2a5, one of the major amylases secreted by the pancreas, shows expression changes in depression and stress mouse models but has no direct relationship with AD^[125]^. SULT1C2 is involved in the metabolism of neurotransmitters and hormones. The sulfotransferase family (such as SULT1A3 and SULT4A1) has been clearly associated with neuropsychiatric diseases including AD, Parkinson’s disease, and schizophrenia, suggesting that SULT1C2 may have similar research value^[126,127]^.The downregulation of Aquaporin-1 (AQP1, 0.41↓) differs from classical findings of water homeostasis dysregulation in AD; previous studies have mostly reported elevated AQP1 expression in AD brains closely associated with Aβ deposition. Its downregulation in urine may reflect terminal depletion of aquaporins from the peripheral circulation^[128,129]^. Vitronectin (VTN, 0.48↓), a multifunctional glycoprotein present in the extracellular matrix, shows downregulation consistent with the pathological feature of extracellular matrix breakdown in AD—VTN participates in cell adhesion, neurogenesis, and blood-brain barrier protection, and is associated with AD and other neurodegenerative diseases^[130]^.

**Table 8.** DEPS Identified at Gestational Day 13 in the Alzheimer’s Disease Model.

| ProteinAccessions | ProteinDescriptions | FC | Pvalue | Reference |
| --- | --- | --- | --- | --- |
| O88839 | Disintegrin and metalloproteinase domain-containing protein 15 | ∞↑ | 3.42E-02 | {117,118,119} |
| P69917 | Translational regulator CsrA | ∞↑ | 4.72E-02 | |
| P9WPE7 | Chaperonin GroEL 2 | ∞↑ | 3.47E-02 | {124} |
| Q6P9J9 | Anoctamin-6 | 121.80 ↑ | 1.36E-02 | {120} |
| Q8BFY6 | Peflin | 50.72 ↑ | 4.91E-02 |  |
| Q8K4G1 | Latent-transforming growth factor beta-binding protein 4 | 25.19 ↑ | 3.23E-02 |  |
| Q6P069 | Sorcin | 24.91 ↑ | 3.14E-02 | {121} |
| Q8VDD5 | Myosin-9 | 11.63 ↑ | 4.42E-02 |  |
| Q9D7S9 | Charged multivesicular body protein 5 | 8.13 ↑ | 1.49E-02 | {123} |
| Q64487 | Receptor-type tyrosine-protein phosphatase delta | 5.53 ↑ | 3.51E-02 |  |
| Q3U0V1;Q92945 | Far upstream element-binding protein 2 | 5.06 ↑ | 3.24E-02 |  |
| Q91VI7 | Ribonuclease inhibitor | 4.62 ↑ | 3.95E-02 |  |
| Q8C7U7 | Polypeptide N-acetylgalactosaminyltransferase 6 | 4.55 ↑ | 4.77E-02 |  |
| Q9R013 | Cathepsin F | 4.23 ↑ | 2.63E-02 |  |
| O35405 | 5'-3' exonuclease PLD3 | 4.05 ↑ | 3.48E-02 | {122} |
| Q64277 | ADP-ribosyl cyclase/cyclic ADP-ribose hydrolase 2 | 3.45 ↑ | 4.49E-02 |  |
| Q61508 | Extracellular matrix protein 1 | 2.59 ↑ | 2.33E-02 |  |
| Q5E924 | Glyceraldehyde-3-phosphate dehydrogenase<br>GAPCP2, chloroplastic | 2.09 ↑ | 2.33E-02 |  |
| Q8R4G6 | Alpha-1,6-mannosylglycoprotein<br>6-beta-N-acetylglucosaminyltransferase A | 0.56 ↓ | 3.70E-02 |  |
| P32921 | Tryptophan--tRNA ligase, cytoplasmic | 0.52 ↓ | 2.55E-02 |  |
| P29788 | Vitronectin | 0.48 ↓ | 1.57E-02 | {131} |
| Q02013 | Aquaporin-1 | 0.41 ↓ | 1.09E-02 | {129,130} |
| P21570 | Angiogenin | 0.34 ↓ | 1.49E-02 |  |
| Q61598 | Rab GDP dissociation inhibitor beta | 0.26 ↓ | 3.10E-02 | {125} |
| P56389 | Cytidine deaminase | 0.17 ↓ | 2.54E-02 |  |
| Q99L13 | 3-hydroxyisobutyrate dehydrogenase, mitochondrial | 0.15 ↓ | 4.09E-02 |  |
| Q9D939 | Sulfotransferase 1C2 | 0.05 ↓ | 3.53E-02 | {127,128} |
| P00688 | Pancreatic alpha-amylase 2a5 | 0.00 ↓ | 3.17E-02 | {126} |

Ras-related protein Rab-35 (∞↑) was the most prominent pathological signal on day 15. Rab35 is a member of the Rab family GTPases, mediating endocytic recycling, actin bundling, and synaptic vesicle turnover. Brain-specific Rab35 knockout mice exhibit anxiety-related behaviors and spatial memory deficits, and Rab35 is essential for the precise positioning of hippocampal pyramidal neurons. Its upregulation in AD patient brains may reflect compensatory stress of the endocytic-recyclingpathway^[131,132]^.Carbonicanhydrase1(CA1,14.83↑)wassignificantly elevated, and carbonic anhydrase may serve as a target for protecting neurovascular function in AD^[133]^. Chloride intracellular channel protein 5 (CLIC5, 8.37↑) showed sustained elevation, suggesting that chloride homeostasis and cytoskeletal connection systems were under high stress^[134]^. Heat shock protein HSP 90-beta (8.20↑), a key chaperone maintaining protein homeostasis, supports the folding and maturation of proteins essential for synaptic transmission and neuronal survival in the nervous system; its upregulation suggests cellular responses to proteotoxic stress^[135]^. The upregulation of Coagulation factor XII (6.79↑) points to AD vascular pathology—Aβ can activate coagulation factor XII, leading to thrombin and bradykinin generation, promoting vascular inflammation and blood-brain barrier damage^[136]^.However, the extreme activation failed to prevent systemic collapse: Spermidine synthase (0.00↓) completely disappeared, with polyamine metabolism exhibiting chronic maladaptive stress responses in AD brains^[137]^; Actin-related protein 2 (0.02↓) marked the collapse of Arp2/3 complex-mediated actin nucleation^[138]^; the sustained downregulation of Rho GDP-dissociation inhibitor 2 (0.22↓) has important pathological significance. Recent studies have confirmed that ARHGDIB (RhoGDIα) can directly bind to tau protein, and its overexpression significantly inhibits tau hyperphosphorylation, delaying pathological progression in AD and vascular dementia models. Its downregulation relieves the inhibition of Rho GTPases, leading to aberrant Rho signaling activation, which in turn exacerbates cytoskeletal disorganization and synaptic damage. Through machine learning combined with weighted gene co-expression network analysis, ARHGDIB has been identified as one of the key genes distinguishing different Braak stages, and its expression changes in microglia are closely associated with AD pathological progression^[139,140]^. Glucose-6-phosphate isomerase (0.10↓) is a tau-related biomarker associated with AD neurodegeneration and cognitive impairment^[141]^. Downregulation of Interleukin-6 receptor subunit alpha (0.32↓) has dual pathological significance—Mendelian randomization analysis indicated that elevated IL-6 receptor levels increase AD risk(OR=1.064, p= 0.003)^[142]^.

**Table 9.**
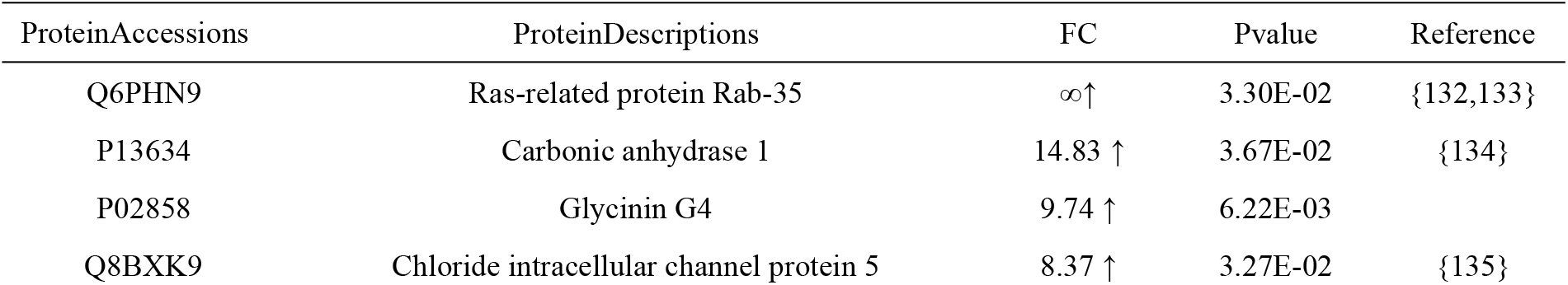
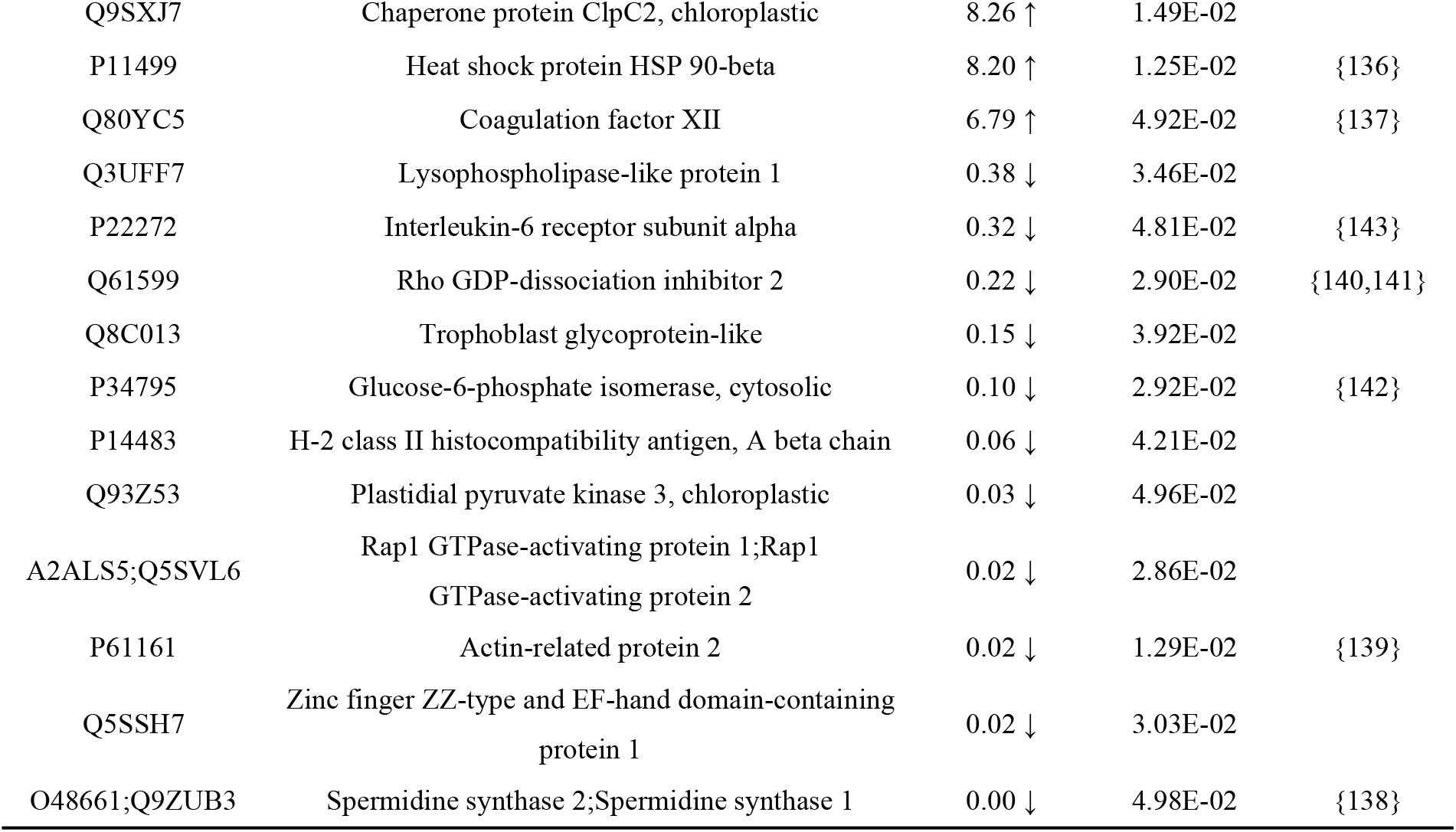
DEPS Identified at Gestational Day 15 in the Alzheimer’s Disease Model.

On day 17, the urinary proteome exhibited a striking “rebound-like multi-pathway extreme activation” phenomenon, with both the number of differential proteins and the magnitude of upregulation greatly increased compared to earlier stages (days 7–15). Multiple pathways simultaneously entered infinite upregulation states, including Eukaryotic translation initiation factor 3 subunit D (EIF3D, ∞↑), Mucin-13 (MUC13, ∞↑), Di-N-acetylchitobiase (CTBS, ∞↑), Keratin, type I cytoskeletal 20 (KRT20, ∞↑), and Thioredoxin reductase 2 (TXNRD2, ∞↑). EIF3D is a key subunit of the eukaryotic translation initiation factor 3 complex. The ExonSkipAD database shows that EIF3D exhibits significant transcript-level differential expression and exon skipping events in multiple brain regions (PCC, CB, TC) of AD patients^[143]^. In microglia of APP/PS1 mouse models, the expression of EIF3 subunits (EIF3H, EIF3E, EIF3K) is significantly upregulated with aging and AD progression^[144]^. MUC13, as a transmembrane mucin predominantly expressed in epithelial tissues, may reflect severe systemic epithelial barrier function stress in the AD model during late gestation^[145]^. Di-N-acetylchitobiase (CTBS, ∞↑) is a member of the chitinase family and participates in glycoprotein degradation. Chitotriosidase-1 (CHIT1, 7.32↑) is a chitinase family protein secreted by activated macrophages, and its upregulation has clear AD pathological evidence. Multiple studies have shown that CSF CHIT1 levels are significantly elevated in AD patients (approximately 133% increase), serving as an important biomarker reflecting microglial activation and neuroinflammation in the brain. Recent plasma proteomic studies have also listed CHIT1 as one of the neuroinflammatory markers significantly associated with Aβ PET status. The sustained elevation of CHIT1 suggests that neuroinflammation remains highly active in AD model mice during late gestation^[146,147]^.Thioredoxin reductase 2, mitochondrial (TXNRD2) is a core enzyme of the mitochondrial thioredoxin system, responsible for clearing mitochondrial reactive oxygen species and maintaining redox homeostasis^[148]^. Studies in Caenorhabditis elegans models have confirmed that TXNRD2 overexpression significantly reduces β-amyloid peptides and amyloid deposits, while its downregulation exacerbates paralysis phenotypes, suggesting a protective role for TXNRD2 against Aβ toxicity^[149]^. KRT20 encodes keratin 20, which is involved in maintaining epithelial cell structural stability, but clear evidence for its differential expression in AD brain tissue is lacking.The simultaneous extreme activation of the above multiple pathways reflects the “terminal struggle” in which the organism mobilizes all compensatory reserves following the superposition of late-gestation physiological burden and AD pathology. Among these, EIF3D-mediated translational regulation and TXNRD2-mediated mitochondrial antioxidant defense have clear AD pathological associations. The coordinated upregulation of Delta-like protein 1 (DLL1, 4.00↑) and Neurogenic locus notch homolog protein 2 (NOTCH2, 3.29↑) further suggests activation of the Notch signaling pathway, which has been shown to participate in AD pathogenesis through regulation of neuroinflammation, synaptic plasticity, and neurogenesis^[150,151]^.The extreme downregulation of TBC1 domain family member 10A (TBC1D10A) has important pathological significance. Proteomic interactome studies using TurboID proximity labeling combined with mass spectrometry have definitively identified TBC1D10A as a core member of the BIN1 interactome—BIN1 being the second largest genetic risk factor for AD. BIN1 directly binds tau through its SH3 domain, participating in synaptic vesicle endocytosis, tau phosphorylation, and pathological tau cell-to-cell transmission, and its dysfunction is a key link in AD pathogenesis. As a component of the BIN1 interaction network, the extreme downregulation of TBC1D10A in AD model urine (from ∞↑ at day 5 to 0.02↓ at day 17) may reflect the collapse and exhaustion of the BIN1-mediated endocytic-recycling negative regulatory system, consistent with the pathological features of BIN1 functional loss in AD brains^[152]^. The extreme downregulation of Selenocysteine lyase (SCLY, 0.03↓) portends severe disruption of selenium metabolism and selenoprotein synthesis. Selenoproteins (such as GPX4 and SELENOM) play irreplaceable roles in maintaining brain redox balance and inhibiting Aβ aggregation and tau phosphorylation^[153]^.

**Table 10.**
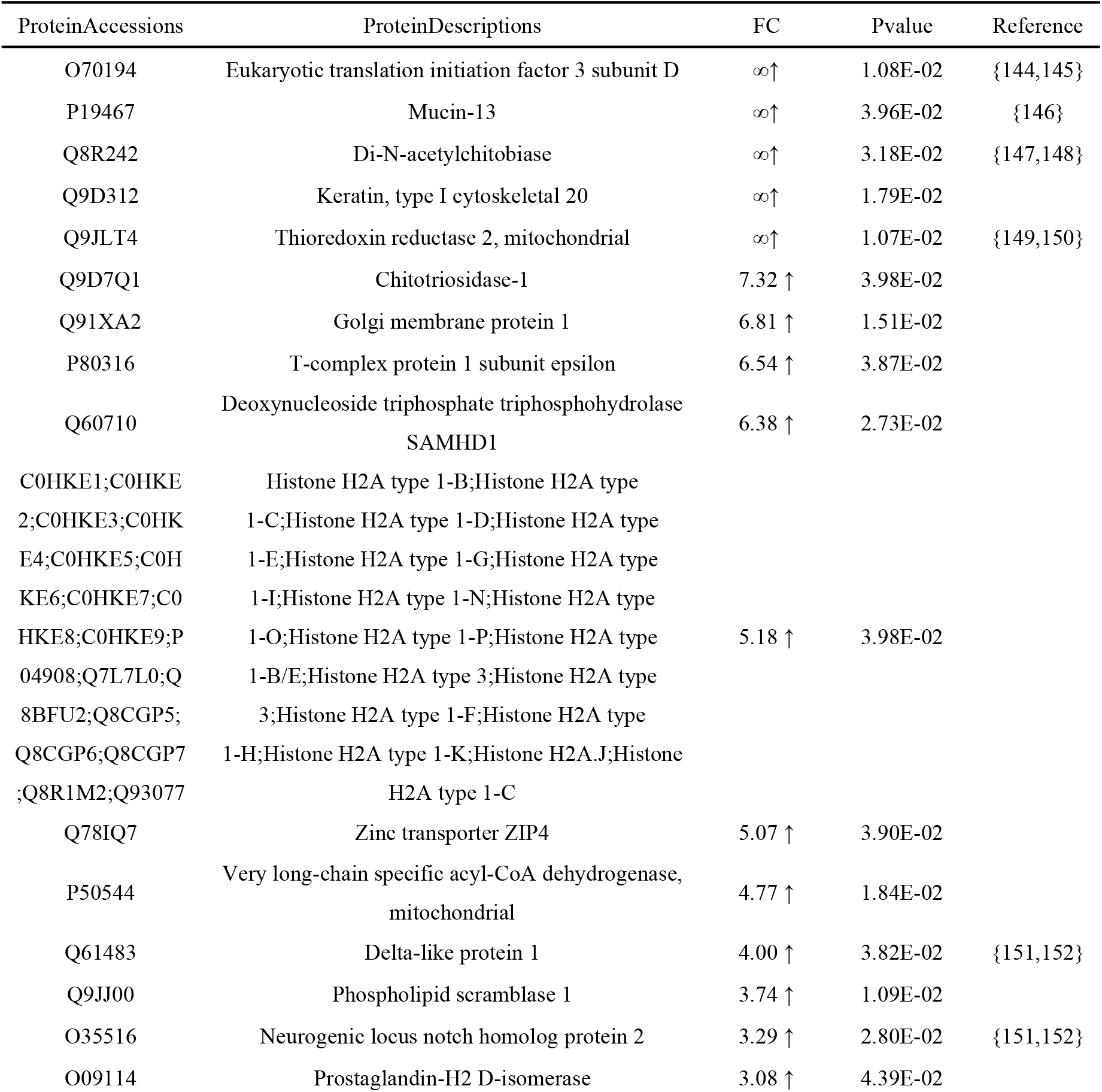
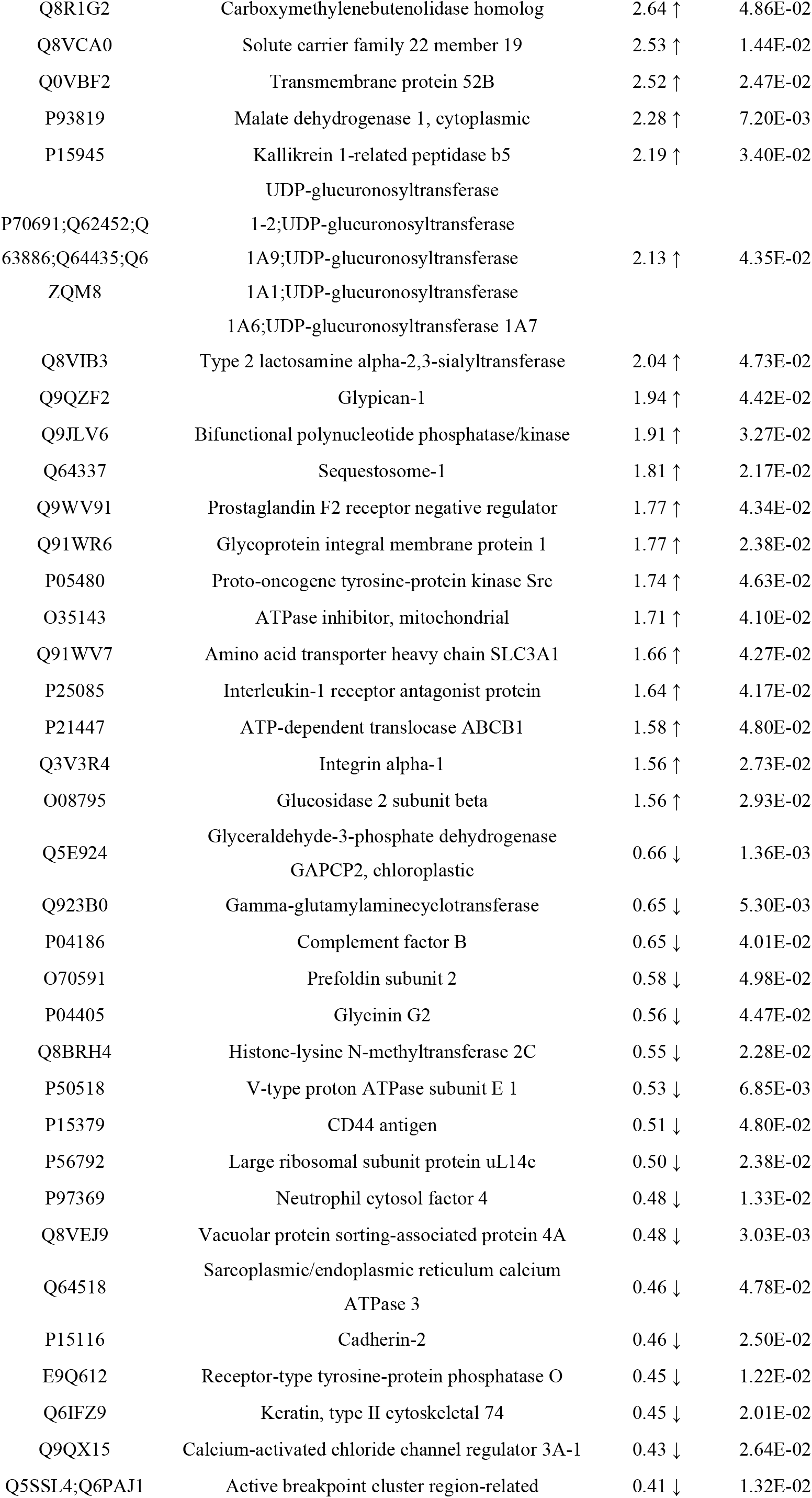
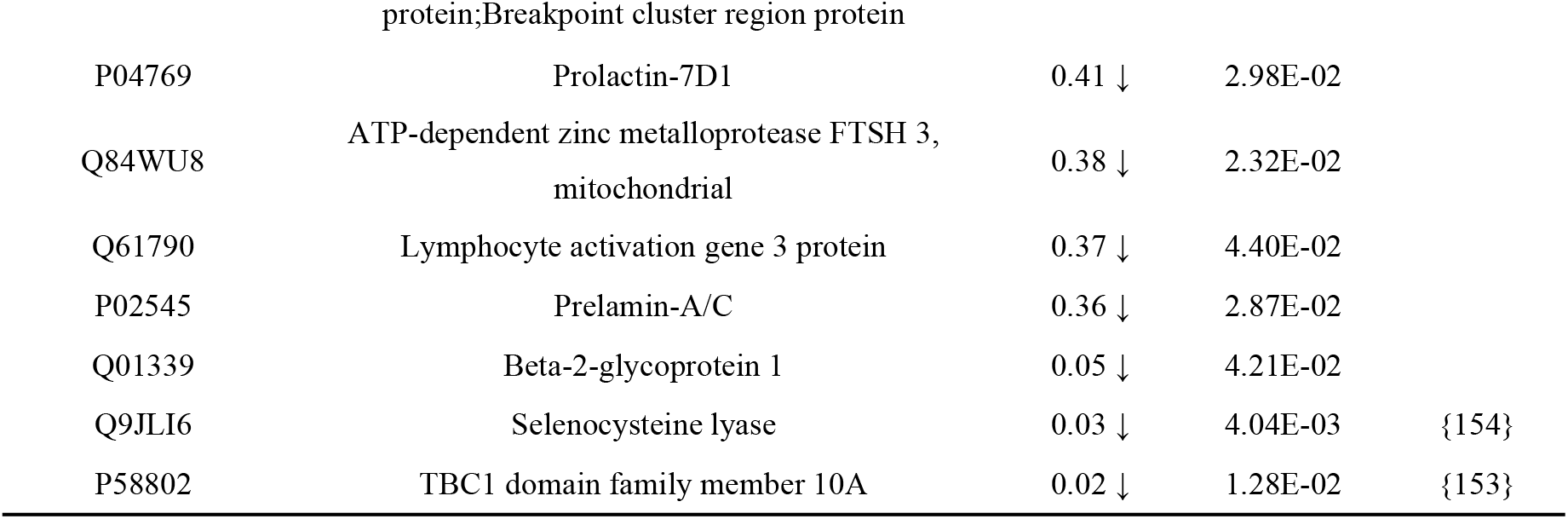
DEPS Identified at Gestational Day 17 in the Alzheimer’s Disease Model.

On day 19, the endpoint of the 19-day gestational course, only 6 differential proteins were identified, forming a stark contrast with the multi-pathway activation of earlier stages in the entire temporal sequence. Near term, maternal mice experience dramatic hormonal fluctuations (oxytocin, cortisol, prostaglandins, etc.) as well as significant physiological changes such as a marked increase in glomerular filtration rate (approximately 50% increase). These labor-related signals may create high-intensity “background noise” in the urinary proteome, diluting or masking non-extreme differential protein signals^[154,155]^. Furthermore, the serum proteomic differences between AD model (5xFAD) and wild-type mice are inherently limited; even with deep coverage detecting approximately 4,000 proteins, the number of differential proteins is modest. Thus, at the peak of physiological stress, only proteins with extremely high abundance or extreme changes (such as 35.86↑, 0.11↓) can exceed the detection threshold^[156]^.Neuroendocrine protein 7B2 (SCG5, 35.86↑) is a secreted protein widely expressed in neurons and endocrine tissues. In vitro experiments have demonstrated its efficient inhibition of Aβ1-42, Aβ1-40, and α-synuclein fibrillization and aggregate formation at a molar ratio of 1:10. Recombinant 7B2 or adenovirus-mediated overexpression of 7B2 blocked Aβ1-42 neurotoxicity and significantly increased cell survival, while knockdown of 7B2 enhanced Aβ1-42-induced cytotoxicity. In APP/PSEN1 mouse brains and in the hippocampus and substantia nigra of human AD and PD patients, 7B2 highly co-localized with Aβ plaques and α-synuclein deposits^[157]^.The upregulation of V-type proton ATPase catalytic subunit A (ATP6V1A, 2.36↑) points to compensatory stress of the lysosomal acidification pathway. ATP6V1A is the catalytic subunit of the V-ATPase proton pump, essential for maintaining hydrolase activity and protein homeostasis. Functional studies have shown that ATP6V1A deficiency affects neuronal process elongation and stability, impairs excitatory synaptic function, and prevents synaptic rearrangement. The underlying mechanisms involve impaired lysosomal pH regulation and autophagic flux obstruction, leading to accumulation of aberrant lysosomes and autophagic vesicles in neuronal cell bodies and synaptic terminals. Its functional loss is closely associated with synaptic dysfunction in neurodevelopmental and neurodegenerative diseases^[158]^. However, Vacuolar protein sorting-associated protein VTA1 homolog (VTA1, 0.11↓), a core component of the ESCRT complex, participates with VPS4 in multivesicular body formation and membrane fission. ESCRT system dysfunction has been confirmed as an important mechanism of failed plasma membrane damage repair in AD and other diseases ^[159,160]^. In addition, Calcium-dependent protein kinase 21 (CDPK21, ∞↑), although clearly annotated in databases such as UniProt as a plant-derived calcium-dependent protein kinase (from rice), has not yet been found to have an association with AD in the literature.

**Table 11.** DEPS Identified at Gestational Day 19 in the Alzheimer’s Disease Model.

| ProteinAccessions | ProteinDescriptions | FC | Pvalue | Reference |
| --- | --- | --- | --- | --- |
| Q9ZSA2 | Calcium-dependent protein kinase 21 | ∞↑ | 4.42E-02 | {162,163} |
| P12961 | Neuroendocrine protein 7B2 | 35.86 ↑ | 4.92E-02 | {158} |
| Q9JJU8 | Adapter Sh3bgrl | 9.13 ↑ | 3.03E-02 |  |
| O08832 | Polypeptide N-acetylgalactosaminyltransferase 4 | 9.08 ↑ | 2.48E-02 |  |
| P50516 | V-type proton ATPase catalytic subunit A | 2.36 ↑ | 3.19E-02 | {159} |
| Q9CR26 | Vacuolar protein sorting-associated protein VTA1 homolog | 0.11 ↓ | 3.68E-02 | {160,161} |

### 3.5 Biological Pathway Analysis of Differential Proteins

GO functional enrichment analysis was performed on the differential proteins at each of the 10 time points using the DAVID database.

GO functional enrichment analysis of the urinary differential proteins at day 1 revealed that multiple biological pathways, including endocytic transport, cytoskeletal dynamic regulation, Aβ metabolism, ESCRT-mediated multivesicular body sorting, and Ras signal transduction, were consistently enriched with high significance (-log10(p-value) ≈ 3.8), suggesting that AD-related pathological changes trigger widespread and coordinated systemic molecular responses as early as early gestation. The enrichment of endocytic transport-related pathways (including endosomal transport, endocytosis, late endosome to lysosome transport, autophagosome maturation, and regulation of endocytic recycling) was highly consistent with the substantial upregulation of endocytosis-related proteins such as Rab-5A (∞↑) at day 1. Studies have confirmed that Rab5A is a core regulator of early endocytic transport, and its aberrant overactivation is an early driver of endocytic dysfunction in AD patients and model mice, leading to early endosome enlargement and accelerated amyloidogenic processing of APP^[161,162]^. The enrichment of ESCRT and multivesicular body assembly-related pathways (multivesicular body assembly, multivesicular body sorting pathway, ubiquitin-dependent protein catabolic process via the multivesicular body sorting pathway) was consistent with VPS4A (5.44↑) upregulation. ESCRT complexes mediate APP sorting into intraluminal vesicles of multivesicular bodies, directing APP toward lysosomal degradation, and ESCRT component deficiency leads to significant intracellular Aβ accumulation^[163]^. The enrichment of cytoskeletal dynamic regulation pathways (regulation of actin cytoskeleton organization, barbed-end actin filament capping) was consistent with F-actin-capping protein subunit beta (4.62↑) upregulation, and F-actin dynamic regulation plays a central role in AD synaptic function and memory maintenance. The enrichment of amyloid-beta clearance by transcytosis and amyloid fibril formation indicated that Aβ metabolism-related pathways had already responded to AD pathological stress at day 1. The enrichment of Ras protein signal transduction and ERK1 and ERK2 cascade was consistent with the widespread upregulation of Ras family members including HRas (3.68↑), R-Ras2 (9.62↑), and Ral-B (13.81↑). The enrichment of defense response to Gram-positive bacterium and positive regulation of immune response suggested early activation of the innate immune system, consistent with CD14 (8.07↑) upregulation. The synchronous activation of these pathways, at the molecular network level, validated the biological relevance of the day 1 proteomic data to early AD endocytic-lysosomal pathway abnormalities, and revealed the systemic response characteristics of AD-related pathological pathways during early gestation.

**Table 12.**
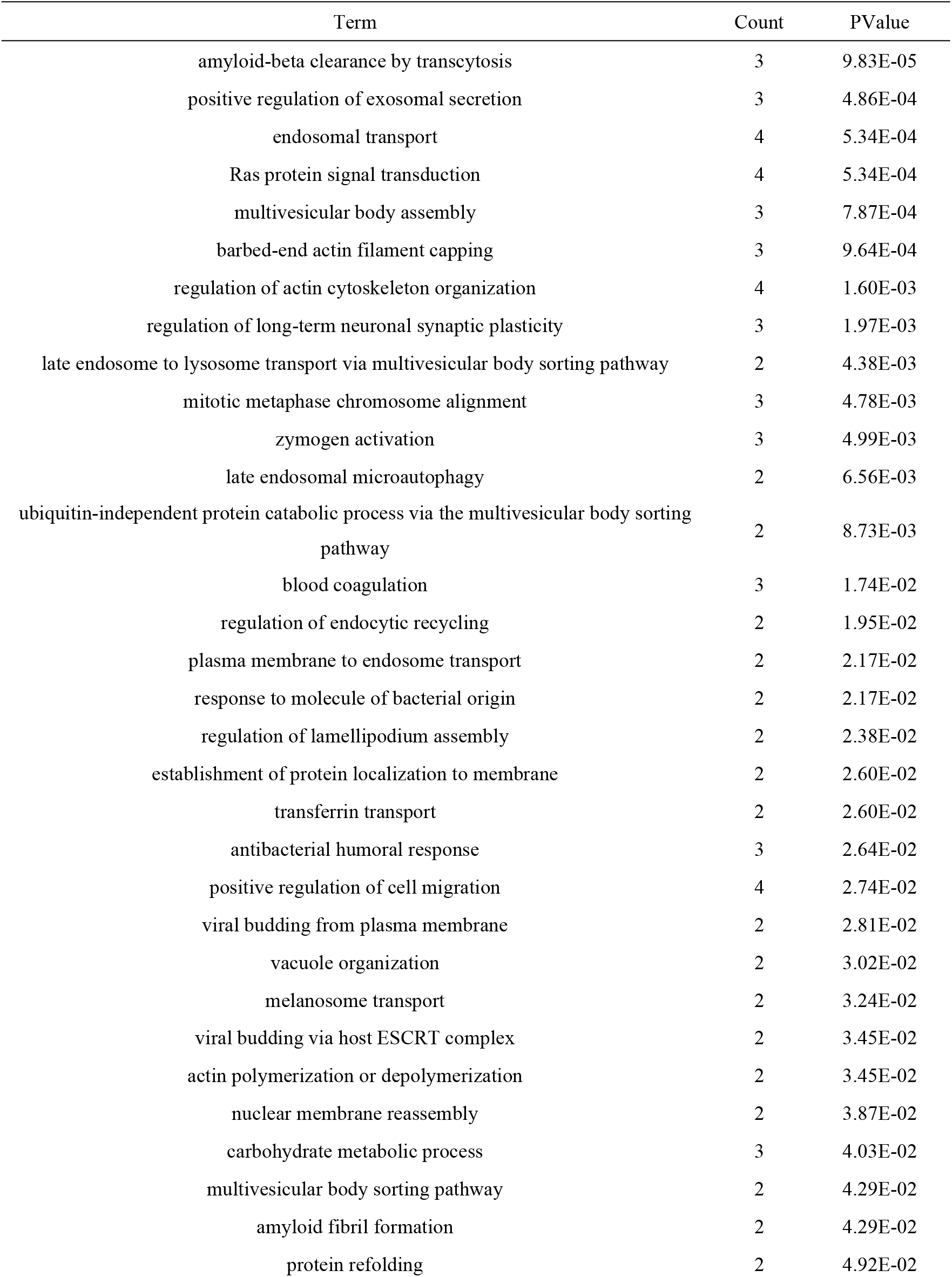
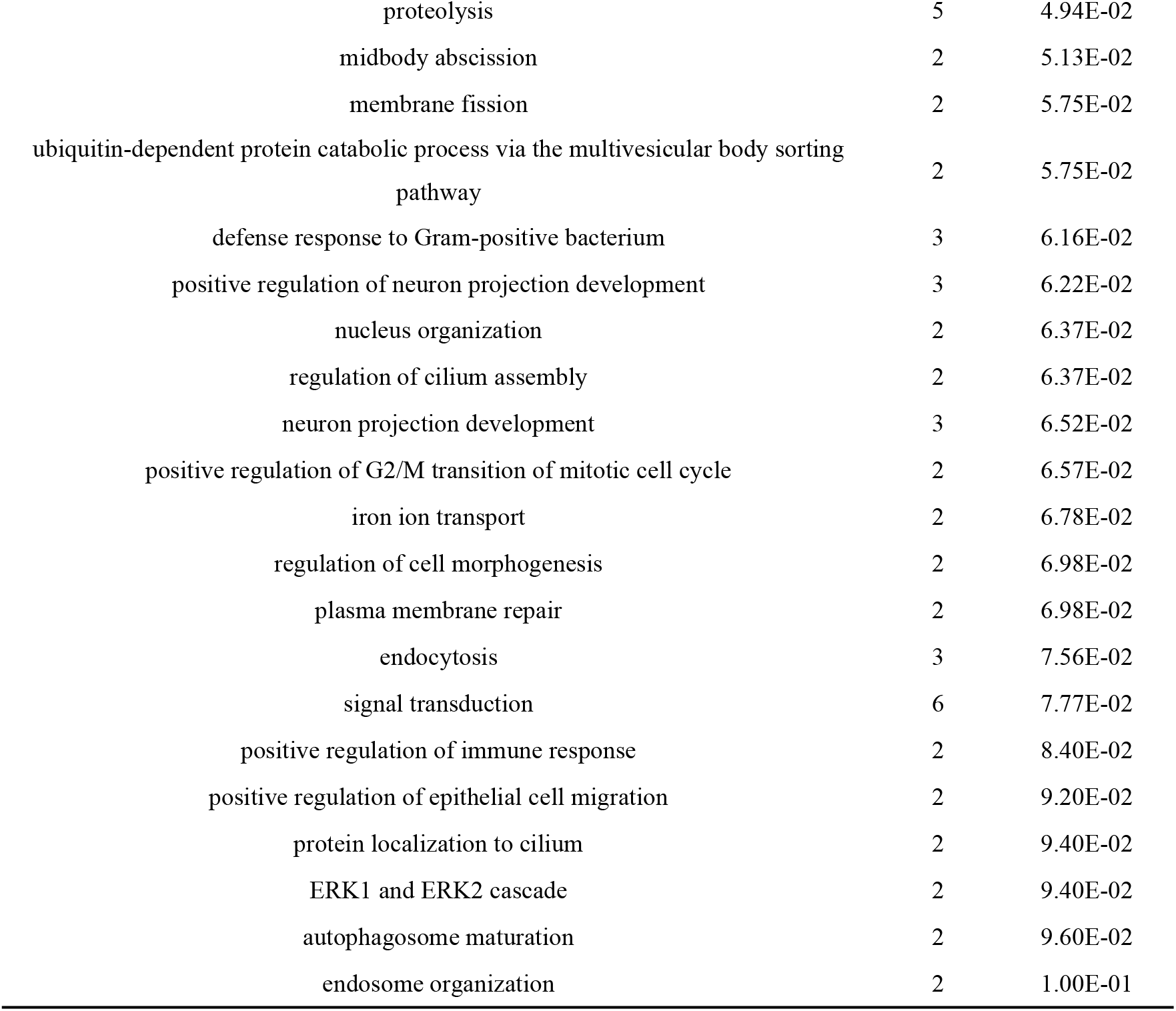
Biological processes enriched by DEPS in the AD model at D1.

| Term | Count | PValue |
| --- | --- | --- |
| amyloid-beta clearance by transcytosis | 3 | 9.83E-05 |
| positive regulation of exosomal secretion | 3 | 4.86E-04 |
| endosomal transport | 4 | 5.34E-04 |
| Ras protein signal transduction | 4 | 5.34E-04 |
| multivesicular body assembly | 3 | 7.87E-04 |
| barbed-end actin filament capping | 3 | 9.64E-04 |
| regulation of actin cytoskeleton organization | 4 | 1.60E-03 |
| regulation of long-term neuronal synaptic plasticity | 3 | 1.97E-03 |
| late endosome to lysosome transport via multivesicular body sorting pathway | 2 | 4.38E-03 |
| mitotic metaphase chromosome alignment | 3 | 4.78E-03 |
| zymogen activation | 3 | 4.99E-03 |
| late endosomal microautophagy | 2 | 6.56E-03 |
| ubiquitin-independent protein catabolic process via the multivesicular body sorting pathway | 2 | 8.73E-03 |
| blood coagulation | 3 | 1.74E-02 |
| regulation of endocytic recycling | 2 | 1.95E-02 |
| plasma membrane to endosome transport | 2 | 2.17E-02 |
| response to molecule of bacterial origin | 2 | 2.17E-02 |
| regulation of lamellipodium assembly | 2 | 2.38E-02 |
| establishment of protein localization to membrane | 2 | 2.60E-02 |
| transferrin transport | 2 | 2.60E-02 |
| antibacterial humoral response | 3 | 2.64E-02 |
| positive regulation of cell migration | 4 | 2.74E-02 |
| viral budding from plasma membrane | 2 | 2.81E-02 |
| vacuole organization | 2 | 3.02E-02 |
| melanosome transport | 2 | 3.24E-02 |
| viral budding via host ESCRT complex | 2 | 3.45E-02 |
| actin polymerization or depolymerization | 2 | 3.45E-02 |
| nuclear membrane reassembly | 2 | 3.87E-02 |
| carbohydrate metabolic process | 3 | 4.03E-02 |
| multivesicular body sorting pathway | 2 | 4.29E-02 |
| amyloid fibril formation | 2 | 4.29E-02 |
| protein refolding | 2 | 4.92E-02 |
| proteolysis | 5 | 4.94E-02 |
| midbody abscission | 2 | 5.13E-02 |
| membrane fission | 2 | 5.75E-02 |
| ubiquitin-dependent protein catabolic process via the multivesicular body sorting pathway | 2 | 5.75E-02 |
| defense response to Gram-positive bacterium | 3 | 6.16E-02 |
| positive regulation of neuron projection development | 3 | 6.22E-02 |
| nucleus organization | 2 | 6.37E-02 |
| regulation of cilium assembly | 2 | 6.37E-02 |
| neuron projection development | 3 | 6.52E-02 |
| positive regulation of G2/M transition of mitotic cell cycle | 2 | 6.57E-02 |
| iron ion transport | 2 | 6.78E-02 |
| regulation of cell morphogenesis | 2 | 6.98E-02 |
| plasma membrane repair | 2 | 6.98E-02 |
| endocytosis | 3 | 7.56E-02 |
| signal transduction | 6 | 7.77E-02 |
| positive regulation of immune response | 2 | 8.40E-02 |
| positive regulation of epithelial cell migration | 2 | 9.20E-02 |
| protein localization to cilium | 2 | 9.40E-02 |
| ERK1 and ERK2 cascade | 2 | 9.40E-02 |
| autophagosome maturation | 2 | 9.60E-02 |
| endosome organization | 2 | 1.00E-01 |

GO functional enrichment analysis of the urinary differential proteins at day 3 revealed that multiple pathways, including endocytic-lysosomal transport, vesicular acidification and protein degradation, chloride homeostasis, and dendritic spine organization, were enriched with high significance (-log10(p-value) approximately 5.0–6.2), suggesting that AD-related pathological changes had progressed from the endocytic initiation stage to the coordinated response level of lysosomal degradation and synaptic function regulation during early gestation. The sustained enrichment of endocytic and vesicular transport pathways (plasma membrane to endosome transport, endosomal transport, endosome to lysosome transport via multivesicular body sorting pathway) was highly consistent with the significant upregulation of endocytosis-related proteins such as Rab-5A (203.22↑) at day 3. Endocytic trafficking abnormalities have been confirmed as drivers of neurodegenerative diseases. Rab5 is a core marker of early endosomes, and its aberrant activation in AD leads to early endosome enlargement and accelerated amyloidogenic processing of APP^[164]^. The enrichment of vacuolar acidification and regulation of autophagosome assembly pointed to lysosomal acidification and autophagic maturation processes. V-ATPase is the core proton pump that maintains the acidic environment of the lysosomal lumen, and V-ATPase dysfunction can impair lysosomal acidification, thereby disrupting the clearance of various substrates (including protein aggregates), and is closely associated with the pathogenesis of AD and other neurodegenerative diseases^[165]^. The enrichment of chloride transport and chloride transmembrane transport suggested that chloride homeostasis regulation was significantly activated at day 3. In AD pathology, Aβ can downregulate KCC2 expression through disruption of the BDNF-TrkB signaling pathway, while abnormally upregulating NKCC1, leading to intracellular chloride accumulation and impaired GABAergic inhibition, thereby promoting neuronal hyperexcitability^[166]^. Studies have confirmed that Aβ1-42 injection leads to significantly elevated NKCC1 expression in the mouse hippocampal CA1 region at 30 days post-injection, suggesting that chloride homeostasis dysregulation is specific to AD^[167]^. The enrichment of dendritic spine organization indicated that the regulation of dendritic spine structure was already affected by AD pathological stress at day 3. Dendritic spines are the postsynaptic sites of most excitatory synapses, and their loss is an early pathophysiological hallmark of AD, which can precede dendritic structural degeneration and overt neurodegeneration, and is closely associated with cognitive dysfunction^[168]^. The maintenance of dendritic spine morphology and density is crucial for synaptic plasticity, and their early impairment in AD is one of the core aspects of synaptic dysfunction^[169]^.

**Table 13.** Biological processes enriched by DEPS in the AD model at D3.

| Term | Count | PValue |
| --- | --- | --- |
| plasma membrane to endosome transport | 3 | 2.32E-04 |
| chloride transport | 4 | 6.19E-04 |
| protein transport | 7 | 1.18E-03 |
| chloride transmembrane transport | 4 | 2.96E-03 |
| intracellular protein transport | 5 | 3.36E-03 |
| angiogenesis | 5 | 4.90E-03 |
| multicellular organism growth | 4 | 7.95E-03 |
| endosomal transport | 3 | 1.20E-02 |
| endocytosis | 4 | 1.22E-02 |
| endosome to lysosome transport via multivesicular body sorting pathway | 2 | 1.37E-02 |
| response to estradiol | 3 | 2.41E-02 |
| dendritic spine organization | 2 | 2.73E-02 |
| phosphate ion transmembrane transport | 2 | 2.95E-02 |
| negative regulation of synaptic transmission | 2 | 3.18E-02 |
| vacuolar acidification | 2 | 3.18E-02 |
| receptor catabolic process | 2 | 3.18E-02 |
| response to growth hormone | 2 | 3.40E-02 |
| epidermal cell differentiation | 2 | 3.62E-02 |
| regulation of autophagosome assembly | 2 | 3.62E-02 |
| cell migration | 4 | 4.12E-02 |
| renal sodium ion absorption | 2 | 4.29E-02 |
| transmembrane transport | 5 | 4.33E-02 |
| vacuolar transport | 2 | 4.51E-02 |
| multivesicular body sorting pathway | 2 | 4.51E-02 |
| positive regulation of monocyte chemotaxis | 2 | 5.17E-02 |
| negative regulation of cell migration | 3 | 5.56E-02 |
| response to cadmium ion | 2 | 5.82E-02 |
| D-glucose import across plasma membrane | 2 | 6.04E-02 |
| response to zinc ion | 2 | 6.47E-02 |
| cellular response to forskolin | 2 | 6.47E-02 |
| negative regulation of gene expression | 4 | 6.79E-02 |
| autophagy | 3 | 7.18E-02 |
| Golgi to plasma membrane transport | 2 | 7.76E-02 |
| extrinsic apoptotic signaling pathway via death domain receptors | 2 | 7.97E-02 |
| protein catabolic process | 3 | 7.98E-02 |
| regulation of endocytosis | 2 | 9.03E-02 |
| ephrin receptor signaling pathway | 2 | 9.45E-02 |
| potassium ion import across plasma membrane | 2 | 9.66E-02 |
| intracellular signal transduction | 4 | 9.75E-02 |

GO functional enrichment analysis of the urinary differential proteins at day 5 revealed that intermediate filament organization and assembly was the most significant core biological process (Count=13, P=2.80E-21), with its enrichment significance far exceeding that of other terms, suggesting that the cytoskeletal system was undergoing profound pathological remodeling at day 5. Intermediate filaments are core components of the neuronal cytoskeleton, including neurofilaments and keratins, and their pathological significance in AD has been widely documented. Neurofilaments constitute the core components of paired helical filaments (PHFs)—the hallmark pathological structures of AD—and PHFs are the basic structural units of neurofibrillary tangles^[170,171]^. Intermediate filament proteins are closely associated with ubiquitination modifications; in AD neurofibrillary tangles, neurofilaments co-localize with ubiquitin, suggesting that cells clear pathogenic protein aggregates through both the ubiquitin-proteasome and autophagy-lysosomal pathways^[172]^. The enrichment of keratin is particularly noteworthy—previous studies have confirmed that keratin 9 is aberrantly elevated in the cerebrospinal fluid and blood of AD patients, with concentrations significantly higher than those in healthy controls, and keratin 9 has been incorporated into a diagnostic panel with an accuracy of 87% for AD^[173]^. In addition, the enrichment of wound healing-related pathways (P = 8.40E-03) provides a novel perspective on AD pathology—some scholars have proposed the hypothesis that AD may result from dysregulated brain wound healing processes, mapping the four core features of AD (β-amyloid deposition, neurofibrillary tangles, inflammation, and apoptosis) onto the four stages of wound healing (hemostasis, inflammation, repair, and remodeling)^[174]^. Meanwhile, the enrichment of lysosome lumen pH regulation (P=1.18E-02) and lysosome acidification (P = 3.02E-02) pointed to the functional state of the endocytic-lysosomal system. Recent review articles have indicated that the precise maintenance of the acidic microenvironment within the lysosomal lumen is critical for neuronal protein degradation and autophagic flux, and that lysosomal pH dysregulation caused by V-ATPase proton pump dysfunction is a common pathogenic mechanism in AD and other neurodegenerative diseases, impairing autophagic flux and triggering neuroinflammation^[175]^. Another study confirmed that the assembly of V-ATPase subunits depends on acetylated microtubules, and that impaired microtubule stability in AD directly leads to lysosomal acidification defects and Aβ accumulation^[176]^. The synchronous enrichment of the above pathways was highly consistent with the substantial upregulation of multiple keratins (Keratin, type I cytoskeletal 16 at 23.62↑, Keratin, type II cytoskeletal 5 at 4.75↑, etc.), LAMP1 (4.16↑), and CHMP5 (7.73↑) observed at day 5, validating, at the cellular biology level, the biological relevance of the day 5 proteomic data to AD cytoskeletal pathology and compensatory activation of the lysosomal system。

**Table 14.** Biological processes enriched by DEPS in the AD model at D5.

| Term | Count | PValue |
| --- | --- | --- |
| intermediate filament organization | 13 | 2.80E-21 |
| morphogenesis of an epithelium | 7 | 9.75E-10 |
| keratinization | 7 | 4.31E-09 |
| epithelial cell differentiation | 7 | 2.28E-08 |
| skin development | 4 | 2.52E-04 |
| keratinocyte differentiation | 4 | 3.44E-04 |
| sarcomere organization | 3 | 2.91E-03 |
| establishment of skin barrier | 3 | 3.03E-03 |
| adenylate cyclase-modulating G protein-coupled receptor signaling pathway | 3 | 3.96E-03 |
| desmosome maintenance | 2 | 5.09E-03 |
| wound healing | 3 | 8.40E-03 |
| positive regulation of NAD(P)H oxidase activity | 2 | 1.02E-02 |
| lysosomal lumen pH elevation | 2 | 1.18E-02 |
| intermediate filament bundle assembly | 2 | 1.35E-02 |
| desmosome organization | 2 | 1.35E-02 |
| keratinocyte development | 2 | 2.19E-02 |
| positive regulation of urine volume | 2 | 2.52E-02 |
| fatty acid metabolic process | 3 | 2.66E-02 |
| lysosomal lumen acidification | 2 | 3.02E-02 |
| cytoskeleton organization | 3 | 3.24E-02 |
| intermediate filament cytoskeleton organization | 2 | 3.51E-02 |
| nucleus organization | 2 | 4.98E-02 |
| positive regulation of superoxide anion generation | 2 | 5.14E-02 |
| adherens junction organization | 2 | 5.62E-02 |
| cell-cell adhesion | 3 | 6.05E-02 |
| protein stabilization | 3 | 6.42E-02 |
| regulation of endocytosis | 2 | 6.74E-02 |
| cell motility | 2 | 7.06E-02 |
| autophagosome maturation | 2 | 7.53E-02 |
| positive regulation of vascular associated smooth muscle cell proliferation | 2 | 7.69E-02 |
| negative regulation of apoptotic signaling pathway | 2 | 7.85E-02 |
| endosome to lysosome transport | 2 | 8.16E-02 |

At day 7, GO enrichment analysis identified only one significantly enriched term, “immunoglobulin mediated immune response” (Count =5, P =1.26E-05). The specific enrichment of this pathway was highly consistent with the significant upregulation of immunoglobulin-related proteins (Ig kappa chain V-II region at 14.84↑ and Ig gamma-1 chain C region at 3.11↑) among the differential proteins at day 7, suggesting that humoral immune responses were specifically activated on gestational day 7.Immunoglobulin-mediated immune responses play a dual role in AD pathology. Classical studies have confirmed that immunoglobulin-positive neurons in AD brains exhibit degenerative morphology and co-localize with complement C1q and C5b-9. The activated microglia surrounding these neurons are located significantly closer to Ig-positive neurons than to Ig-negative neurons, suggesting that these neurons may be undergoing antibody-dependent classical complement pathway-mediated death^[177]^. In amyloidosis mouse models, inhibiting neuronal firing reduces complement deposition and protects synapses, indicating that immunoglobulin-complement-mediated synaptic pruning plays an important role in early AD pathology. Furthermore, recent studies have confirmed that gamma-type immunoglobulins (IgG) can localize to the brain parenchyma of multiple amyloid pathology mouse models and specifically enrich in microglia rather than other brain cell types. Microglia exposed to IgG exhibit enhanced phagocytic capacity, suggesting that immunoglobulins may exert protective effects by enhancing microglial Aβ clearance^[178]^. The specific enrichment of this pathway was highly consistent with the substantial upregulation of Ig kappa chain and Ig gamma chain proteins at day 7, indicating, at the functional level, that AD pathology had entered a stage of specific activation of immunoglobulin-mediated humoral immune responses on gestational day 7.

**Table 15.** Biological processes enriched by DEPS in the AD model at D7.

| Term | Count | PValue |
| --- | --- | --- |
| immunoglobulin mediated immune response | 5 | 1.26E-05 |

At day 9, GO functional enrichment analysis revealed that “heterochromatin formation” was the most highly enriched term (Count = 18, P = 2.85E-32), indicating that chromatin remodeling and epigenetic regulation constituted the core molecular events of AD pathology at gestational day 9. Heterochromatin loss is one of the important early features of neurodegeneration: Bmi1 heterozygous mice (Bmi1⁺/⁻) exhibit age-dependent heterochromatin loss and activation of DNA damage responses, along with AD-like cognitive deficits and neurodegenerative pathology^[179]^. This study further confirmed that heterochromatin abnormalities and DNA damage responses at repetitive DNA sequences are also present in AD patient brain tissues, suggesting that heterochromatin instability is a common pathological feature of AD^[179]^. Meanwhile, the enrichment of “protein localization to CENP-A containing chromatin” (Count = 2, P = 2.83E-02) suggested perturbation of centromeric chromatin structure. The term “immunoglobulin mediated immune response” was also enriched at day 9 (Count = 3, P = 3.52E-02), forming a temporal continuation with the enrichment results from day 7. Immunoglobulin-positive neurons in AD brains exhibit degenerative morphology and co-localize with complement C1q and C5b-9, with surrounding activated microglia located significantly closer to Ig-positive neurons than to Ig-negative neurons, suggesting that these neurons may be undergoing antibody-dependent classical complement pathway-mediated death [180]. Recent studies have further found that in amyloidosis mouse models, B cells can migrate from the meninges to the brain parenchyma and secrete IgM, which recruits complement C1q to tag synapses for pruning, and inhibiting neuronal firing reduces complement deposition and protects synapses^[181]^. The enrichment of “positive regulation of chemokine (C-X-C motif) ligand 2 production” (Count = 2, P = 2.27E-02) is consistent with the role of chemokines in AD—chemokines can participate in AD progression by disrupting blood-brain barrier integrity, promoting immune cell infiltration, and exacerbating Aβ production and tau hyperphosphorylation^[182]^. The synchronized enrichment of the above pathways was highly consistent with the sharp downregulation of cytoskeletal proteins such as TUBA4A (0.11↓) and extracellular matrix proteins such as THBS1 (0.05↓) observed at day 9. The extreme significance of heterochromatin formation enrichment suggests that AD-related pathological signals had penetrated to the core level of gene expression regulation—epigenetic reprogramming—by day 9 of gestation, which may be the upstream driving mechanism of the widespread protein expression dysregulation observed earlier.

**Table 16.** Biological processes enriched by DEPS in the AD model at D9.

| Term | Count | PValue |
| --- | --- | --- |
| heterochromatin formation | 18 | 2.85E-32 |
| positive regulation of chemokine (C-X-C motif) ligand 2 production | 2 | 2.27E-02 |
| protein localization to CENP-A containing chromatin | 2 | 2.83E-02 |
| immunoglobulin mediated immune response | 3 | 3.52E-02 |
| positive regulation of T cell differentiation | 2 | 6.26E-02 |
| microtubule-based process | 2 | 6.53E-02 |
| positive regulation of double-strand break repair via homologous recombination | 2 | 6.80E-02 |
| chromatin organization | 3 | 7.99E-02 |

At day 11, GO functional enrichment analysis revealed that pathways including maintenance of translational fidelity, presynaptic/postsynaptic translation, ribosomal small subunit biogenesis, and positive regulation of canonical Wnt signaling pathway were consistently enriched with high significance. The extreme enrichment of the maintenance of translational fidelity pathway has profound pathological implications—neurons are particularly sensitive to chronic loss of translational fidelity, and aberrant proteins generated by translation errors can trigger protein aggregation and cytotoxicity. Mutations in multiple translation machinery components (including tRNA synthetases, elongation factors, and ribosomal proteins) have been linked to neurodegenerative diseases^[183]^. The enrichment of presynaptic and postsynaptic translation pathways is closely associated with neurological disorders including fragile X syndrome, autism spectrum disorders, and cognitive impairment^[184,185]^. The enrichment of ribosomal small subunit biogenesis is consistent with recent evidence of ribosomal dysfunction in AD—multiple studies have found that ribosome biogenesis is one of the most significantly impaired biological processes in AD, and ribosomal dysfunction has been proposed as a new pathological mechanism of AD independent of Aβ aggregation and tau phosphorylation^[185,186]^. The enrichment of positive regulation of canonical Wnt signaling pathway indicates that this pathway was significantly activated at day 11—the Wnt/β-catenin signaling pathway is crucial for blood-brain barrier integrity and Aβ clearance, and its dysfunction can promote AD progression. In early AD models, the upregulation of Dkk-1 is closely associated with GSK3β activation and tau hyperphosphorylation[187,188].

**Table 17.** Biological processes enriched by DEPS in the AD model at D11.

| Term | Count | PValue |
| --- | --- | --- |
| maintenance of translational fidelity | 11 | 3.25E-29 |
| translation at presynapse | 11 | 1.67E-20 |
| translation at postsynapse | 11 | 1.97E-20 |
| ribosomal small subunit biogenesis | 11 | 6.21E-19 |
| positive regulation of canonical Wnt signaling pathway | 11 | 2.61E-17 |
| cytoplasmic translation | 11 | 1.14E-16 |
| translation | 11 | 5.10E-14 |
| cell activation | 2 | 1.27E-02 |
| vesicle budding from membrane | 2 | 1.95E-02 |
| establishment of endothelial barrier | 2 | 2.14E-02 |
| lipoprotein metabolic process | 2 | 3.38E-02 |
| regulation of heart rate by cardiac conduction | 2 | 3.76E-02 |
| lipid metabolic process | 3 | 3.82E-02 |
| intermediate filament organization | 2 | 7.11E-02 |
| phagocytosis | 2 | 8.57E-02 |
| cholesterol metabolic process | 2 | 9.64E-02 |

At day 13, GO functional enrichment analysis revealed that “angiogenesis” was the most highly enriched term (Count = 5, P = 1.38E-04), suggesting that dysregulated angiogenesis constituted one of the core molecular events of AD pathology at gestational day 13. Pathological angiogenesis exists in AD patient brains, characterized by an association between cerebrovascular damage and neurodegeneration^[189]^. Single-nucleus transcriptomic studies have further revealed significant abnormalities in angiogenesis-related transcriptional features in AD patient brain endothelial cells, with increased β-amyloid associated with vascular barrier damage and dysregulated angiogenic responses^[190]^. In addition, the enrichment of “plasma membrane repair” (Count = 2, P = 2.95E-02) is consistent with membrane damage mechanisms in AD—recent studies have found that Aβ induces neuronal plasma membrane damage, and intracellular calcium elevation and calpain activation are driving factors for dystrophic neurite formation, with annexin A6-mediated plasma membrane repair reducing dystrophic neurites and phosphorylated tau accumulation^[191]^. The enrichment of “integrin-mediated signaling pathway” (Count = 2, P = 9.51E-02) is consistent with aberrant integrin signaling activation in AD—Aβ mediates neurotoxicity through α2β1 and αVβ1 integrins, and integrin signaling pathway activation of Pyk2 kinase is a key step in Aβ-induced neuronal damage [192]. The enrichment of “negative regulation of translation in response to stress” (Count = 2, P = 9.03E-03) and “tRNA stabilization” (Count = 2, P = 1.26E-02) points to the translational regulation level—tRNA modification deficiency can lead to tRNA instability and protein homeostasis dysregulation, subsequently triggering neurodegeneration^[193]^. The synchronized enrichment of the above pathways was highly consistent with the extreme upregulation of stress-related proteins such as ADAM15 (∞↑) and the severe downregulation of metabolic enzyme systems observed at day 13.

**Table 18.**
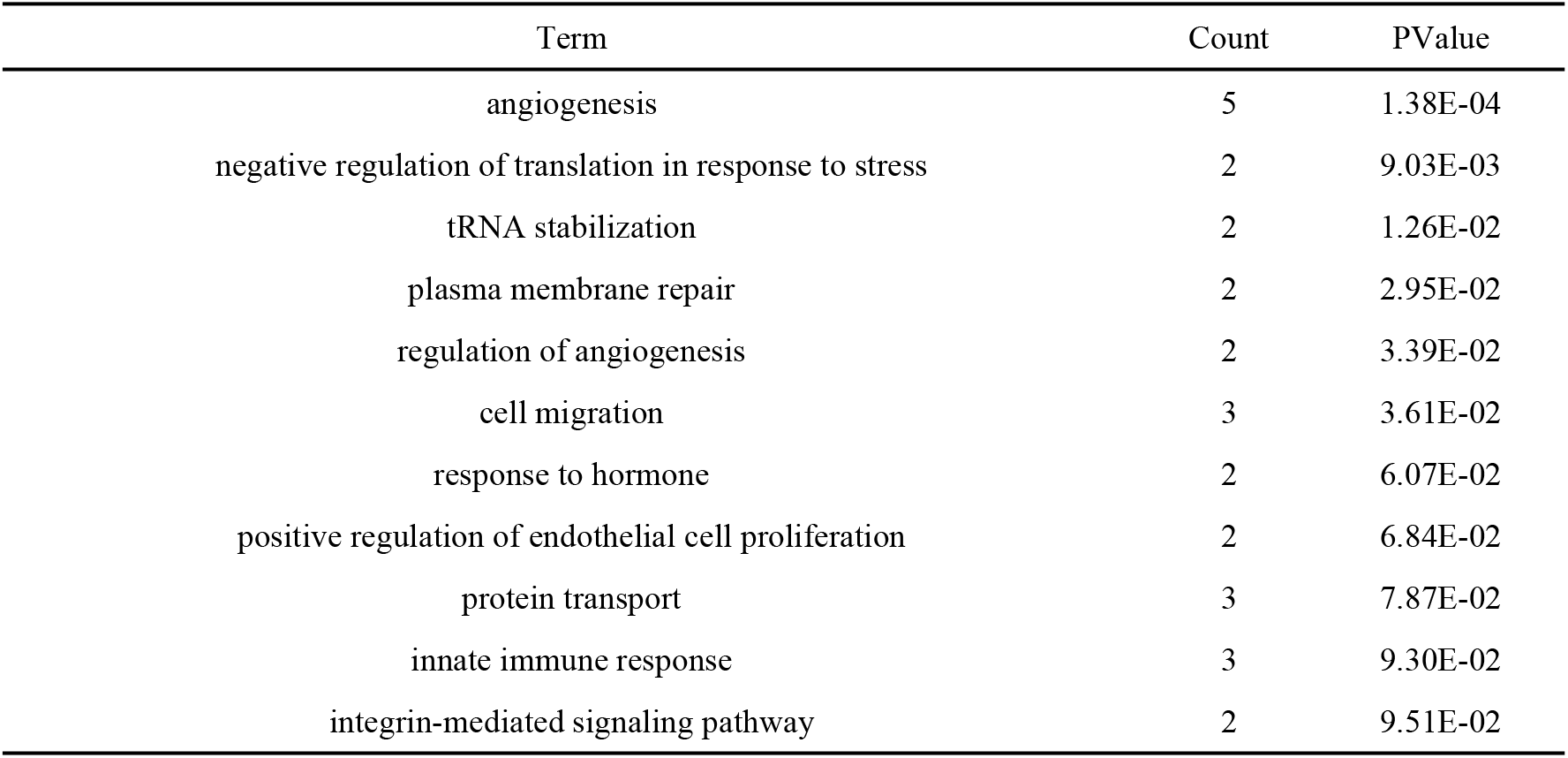
Biological processes enriched by DEPS in the AD model at D13.

| Term | Count | PValue |
| --- | --- | --- |
| angiogenesis | 5 | 1.38E-04 |
| negative regulation of translation in response to stress | 2 | 9.03E-03 |
| tRNA stabilization | 2 | 1.26E-02 |
| plasma membrane repair | 2 | 2.95E-02 |
| regulation of angiogenesis | 2 | 3.39E-02 |
| cell migration | 3 | 3.61E-02 |
| response to hormone | 2 | 6.07E-02 |
| positive regulation of endothelial cell proliferation | 2 | 6.84E-02 |
| protein transport | 3 | 7.87E-02 |
| innate immune response | 3 | 9.30E-02 |
| integrin-mediated signaling pathway | 2 | 9.51E-02 |

At day 15, GO functional enrichment analysis of the urinary differential proteins revealed significant enrichment of pathways including regulation of small GTPase-mediated signal transduction, antigen processing and presentation, and cellular response to type II interferon. The enrichment of small GTPase signal transduction regulation was highly consistent with the sharp downregulation of Rap1 GTPase-activating protein (0.02↓) and ARHGDIB (0.22↓) at day 15. The small GTPase superfamily (Ras, Rho, Rab, etc.) constitutes core molecular switches that regulate cytoskeletal dynamics, membrane trafficking, and signal transduction, with their activity precisely regulated by three classes of factors: GEFs, GAPs, and GDIs^[194]^. The RhoA/ROCK/GSK3β signaling axis participates in AD progression by promoting tau hyperphosphorylation, synaptic damage, and neuroinflammation^[195]^. Small GTPase regulatory abnormalities have been confirmed as key links in autophagy, lysosomal function, and tau aggregation in AD. The enrichment of antigen processing and presentation indicated that adaptive immunity was significantly activated at day 15^[194]^. In AD, microglial MHC II-dependent antigen presentation is enhanced, accompanied by T cell infiltration, which is a key driver of APOE4-related neuroinflammation^[196].^ HLA/MHC gene loci are among the most significant genetic risk factors for AD, with HLA gene variants significantly associated with AD risk^[197,198]^. The enrichment of cellular response to type II interferon is consistent with T cell infiltration and IFN-γ signaling activation in AD. The choroid plexus of J20 AD mouse models shows significant expression changes in type II interferon response genes^[199]^. IFN-γ signaling can induce microglial polarization toward a protective phenotype, enhancing amyloid plaque compaction and reducing dystrophic neurites^[200]^. The synchronized enrichment of the above pathways suggests that AD-related pathological signals had extended from the earlier (days 1–13) endocytic-lysosomal stress and cytoskeletal remodeling to the deeper coordinated level of small GTPase signaling network collapse and adaptive immune activation by gestational day 15.

**Table 19.** Biological processes enriched by DEPS in the AD model at D15.

| Term | Count | PValue |
| --- | --- | --- |
| regulation of small GTPase mediated signal transduction | 2 | 0.01463114 |
| antigen processing and presentation | 2 | 0.02090972 |
| cellular response to type II interferon | 2 | 0.048474504 |

At day 17, GO functional enrichment analysis revealed that “heterochromatin formation” was the most highly enriched core term (Count = 18, P = 2.03E-26), while metabolic pathways including “flavone metabolic process,” “sterol metabolic process,” “estrogen metabolic process,” and “cellular response to glucocorticoid stimulus” were also significantly enriched, suggesting that AD pathology had progressed to the coordinated response level of chromatin remodeling, lipid metabolic reprogramming, and hormonal signaling by gestational day 17. Histone post-translational modifications and heterochromatin structural changes have been confirmed as core pathological events in AD—heterochromatin loss leads to increased genomic instability, transposon activation, and DNA damage accumulation, constituting an important driving mechanism of neurodegeneration^[201]^. Heterochromatin formation was also observed at day 9. The enrichment of sterol metabolism and estrogen metabolism pathways is consistent with lipid homeostasis dysregulation in AD—sterol metabolism plays a central role in maintaining neuronal membrane integrity and myelin formation, while estrogen, as a key regulator of bioenergetic systems, can exacerbate mitochondrial dysfunction and Aβ pathology when its metabolism is abnormal^[202]^. The enrichment of flavone metabolic pathways is consistent with the neuroprotective effects of flavonoids in AD—epidemiological studies have shown that long-term dietary intake of flavonoid-rich foods is associated with reduced AD risk, and flavonoids exert neuroprotective effects through multiple mechanisms including antioxidant, anti-inflammatory, and Aβ metabolism regulation ^[203,204]^. The synchronized enrichment of the above pathways was highly consistent with the extreme activation of multiple pathways including EIF3D (∞↑) and MUC13 (∞↑), and the collapse-level downregulation of key regulatory proteins such as TBC1D10A (0.02↓) observed at day 17, suggesting that AD-related pathological signals had extended from the earlier (days1–15) translational regulation and signal transduction abnormalities to systemic response level of chromatin remodeling, lipid metabolic reprogramming, and hormonal signaling networks by gestational day 17.

**Table 20.** Biological processes enriched by DEPS in the AD model at D17.

| Term | Count | PValue |
| --- | --- | --- |
| heterochromatin formation | 18 | 2.03E-26 |
| flavone metabolic process | 5 | 3.90E-09 |
| sterol metabolic process | 5 | 4.78E-07 |
| liver development | 7 | 1.10E-06 |
| estrogen metabolic process | 5 | 4.29E-06 |
| cellular response to glucocorticoid stimulus | 5 | 4.29E-06 |
| negative regulation of respiratory burst | 2 | 5.58E-03 |
| negative regulation of blood vessel remodeling | 2 | 8.36E-03 |
| negative regulation of neutrophil degranulation | 2 | 8.36E-03 |
| negative regulation of cellular extravasation | 2 | 8.36E-03 |
| placenta development | 3 | 1.31E-02 |
| negative regulation of macrophage migration | 2 | 1.39E-02 |
| cell migration | 5 | 1.40E-02 |
| proximal tubule development | 2 | 1.67E-02 |
| inner ear morphogenesis | 3 | 1.87E-02 |
| negative regulation of regulatory T cell differentiation | 2 | 1.94E-02 |
| macrophage migration | 2 | 2.21E-02 |
| neutrophil degranulation | 2 | 2.21E-02 |
| protein K63-linked ubiquitination | 3 | 2.22E-02 |
| chitin catabolic process | 2 | 2.49E-02 |
| phagocytosis | 3 | 2.69E-02 |
| marginal zone B cell differentiation | 2 | 2.76E-02 |
| regulation of vascular permeability | 2 | 3.57E-02 |
| left/right axis specification | 2 | 3.84E-02 |
| placenta blood vessel development | 2 | 4.11E-02 |
| epithelial cell development | 2 | 4.65E-02 |
| type B pancreatic cell development | 2 | 4.65E-02 |
| protein localization to CENP-A containing chromatin | 2 | 5.44E-02 |
| carbohydrate metabolic process | 3 | 6.23E-02 |
| negative regulation of inflammatory response | 3 | 6.30E-02 |
| positive regulation of Ras protein signal transduction | 2 | 6.76E-02 |
| modulation of chemical synaptic transmission | 3 | 7.00E-02 |
| positive regulation of protein localization | 2 | 7.28E-02 |
| DNA damage response | 4 | 7.53E-02 |
| neuronal stem cell population maintenance | 2 | 7.54E-02 |
| negative regulation of cell migration | 3 | 7.79E-02 |
| nucleus organization | 2 | 8.06E-02 |
| temperature homeostasis | 2 | 8.06E-02 |
| negative regulation of endothelial cell proliferation | 2 | 8.57E-02 |
| regulation of Rho protein signal transduction | 2 | 8.83E-02 |
| focal adhesion assembly | 2 | 8.83E-02 |
| activation of GTPase activity | 2 | 9.34E-02 |

Notably, no significantly enriched GO biological pathways were detected at day 19. This result stands in stark contrast to day 17, which showed enrichment of multiple significant terms including heterochromatin formation (Count = 18, P = 2.03E-26), while day 19 yielded only 6 differential proteins belonging to different functional categories that could not form a statistically significant coordinated functional network.

## 4. Discussion

The core innovation of this study lies in the first proposal and validation of the hypothesis that “fetal AD susceptibility status can be non-invasively identified through maternal urine during pregnancy.” Although previous studies have revealed the embryonic developmental origins of AD, they have consistently failed to solve the problem of non-invasive identification of fetal AD-related pathological changes, leaving the source prevention of AD in a long-standing void. Through temporal urinary proteomic analysis covering the entire course of pregnancy, this study systematically demonstrated that fetuses carrying AD susceptibility genes can induce regular, identifiable temporal dynamic changes in the maternal urinary proteome at various stages of embryonic development, providing direct experimental evidence for non-invasive prenatal early warning of AD. One of the most critical findings of this study is the molecular response at the zygote stage. At gestational D1 (corresponding to the zygote formation and pre-implantation stage), maternal urine already showed infinite upregulation of the endocytic pathway key molecule Rab-5A as the core, with simultaneous enrichment of multiple AD-related pathways including Aβ metabolism, endocytic transport, ESCRT, and Ras signal transduction. At this stage, the embryo’s own protein synthesis capacity is limited, primarily relying on maternal ooplasmic reserves of mRNA and proteins. However, zygotes carrying APP/PS1/Tau mutations already initiate aberrant gene transcription and cellular signaling activation at the zygote stage. Through early maternal-fetal bidirectional communication, abnormal signal molecules released by the zygote can trigger systemic micro-responses in the mother, allowing AD-related molecular changes to be captured by the urinary proteome before tissue phenotype manifestation. This finding substantially moves the time window for AD pathological detection forward to the very beginning of individual development, breaking through the temporal limitations of traditional morphological detection.

The temporal dynamic features during pregnancy provide a high-resolution molecular atlas for understanding the embryonic origin of AD. In early gestation (D1–D5), with Rab5A-mediated endocytic pathway abnormality as the core, the endocytic-lysosomal system and cytoskeletal regulatory pathways were progressively activated, reflecting early compensatory responses to AD-related proteotoxic stress. In mid-gestation (D7–D11), pathological signals extended from the endocytic-lysosomal level to chromatin remodeling (D9, heterochromatin formation enrichment, P = 2.85E-32) and synaptic local translation regulation (D11, maintenance of translational fidelity and pre/postsynaptic translation enrichment), marking the entry of fetal neural developmental defects into an irreversible molecular patterning stage—this is a key node for identifying the clinical intervention window. In late gestation (D13–D17), the differential protein profile exhibited a polarized pattern of “multi-pathway extreme activation coexisting with systemic collapse,” with angiogenesis, small GTPase signaling regulation, steroid metabolism, and interferon response pathways sequentially recruited, reflecting the terminal stage of systemic decompensation following the superposition of AD pathology and late-gestation high metabolic load. At the gestational endpoint D19, the number of differential proteins plummeted to the lowest level of the entire cycle (5 proteins), with no significantly enriched GO pathways, suggesting that the system entered a “response silence” state after experiencing the “last mobilization.” This complete temporal evolution map confirms that maternal urine captures dynamic pathological changes during fetal development rather than fixed genetic background signals, possessing dynamic diagnostic value.

The urinary proteome diagnostic system constructed in this study possesses multiple core advantages. First, the complete non-invasiveness and temporal collectability of urine samples provide natural clinical accessibility in prenatal monitoring scenarios. Second, through the rigorous pairing of the experimental group (3xTg-AD male × wild-type female) and control group (wild-type × wild-type), both groups of maternal dams were of wild-type background, excluding direct disturbance from maternal transgenic proteins and ensuring that all differential molecules and pathways can be specifically attributed to developmental abnormalities mediated by fetal AD pathogenic genes, guaranteeing the specificity of diagnostic features. At the same time, this study has certain limitations. Although the 3xTg mouse model recapitulates core AD pathological features, inherent differences remain between this model and human sporadic AD in terms of etiology and disease course, and the conclusions require further validation in clinical cohorts. The current differential protein identification is based on label-free quantitative mass spectrometry discovery screening and lacks targeted validation (such as PRM or ELISA); subsequent studies should perform independent cohort validation of core candidate proteins.

## 5. Conclusion

Through temporal urinary proteomic analysis covering the entire course of pregnancy, this study for the first time mapped the complete whole-gestation dynamic urinary proteomic profile of maternal mice carrying AD-susceptible gene fetuses, revealing a complete molecular evolution trajectory from zygote to parturition. It demonstrates that maternal gestational urine can serve as a non-invasive, dynamic liquid biopsy sample for prospective identification of fetal AD susceptibility status. This finding provides critical animal experimental foundations for research on the embryonic origin of AD and the clinical translation of non-invasive prenatal monitoring biomarkers.

